# Movement-independent representation of reward-predicting cues in the medial part of the primate premotor cortex

**DOI:** 10.1101/2024.08.24.609512

**Authors:** Keisuke Sehara, Masashi Kondo, Yuka Hirayama, Teppei Ebina, Masafumi Takaji, Akiya Watakabe, Ken-ichi Inoue, Masahiko Takada, Tetsuo Yamamori, Masanori Matsuzaki

## Abstract

Neural activity across the dorsal neocortex of rodents is dominated by orofacial and limb movements, irrespective of whether the movements are task-relevant or task-irrelevant. To examine the extent to which movements and a primitive cognitive signal, i.e., reward expectancy, modulate the activity of multiple cortical areas in primates, we conducted unprecedented wide-field one-photon calcium imaging of frontoparietal and auditory cortices in common marmosets while they performed a classical conditioning task with two auditory cues associated with different reward probabilities. Licking, eye movement, and hand movement strongly modulated the neuronal activity after cue presentation in the motor and somatosensory cortices in accordance with the somatotopy. By contrast, the posterior parietal cortex and primary auditory cortex did not show much influence from licking. Licking increased the activity in the caudal part of the dorsal premotor cortex, but decreased the activity in the central and lateral parts of the rostral part of the dorsal premotor cortex (PMdr). Reward expectancy that was separable from both spontaneous and goal-directed movements was mainly represented in the medial part of PMdr. Our results suggest that the influence of movement on primate cortical activity varies across areas and movement types, and that the premotor cortex processes motor and cognitive information in different ways within further subdivided areas.

## Introduction

Brain activity is ultimately designed to react with the external world in real time through movement. Therefore, it is natural that movement itself modulates brain activity. For example, locomotion largely increases neuronal activity in the mouse primary visual cortex (V1) without changing stimulus selectivity (Niell and Stryker, 2010). Recently, cor-tex-wide calcium imaging and brain-wide electrical recording in mice revealed that a variety of movements dominate the cortical activity not only in the motor cortex and somatosensory cortex, which are directly related to orofacial and body movements, but also in other cortical areas. In particular, orofacial movements such as licking and eye movement significantly increase cortical activity, irrespective of whether such movements are task-relevant or not (Kondo and Matsuzaki, 2021; Musall et al., 2019; Quarta et al., 2022; Salkoff et al., 2020; Stringer et al., 2019; Zagha et al., 2022). However, the effects of movements on the cortical activity of non-human primates remain elusive. The impact of spontaneous facial and body movements on visual cortex (V1, V2, and V3) activity is small, and movement-related change in the activity can be explained by movement-induced shifts in visual input on the retina (Talluri et al., 2023). Running was shown to suppress V1 activity in the common marmoset, but to a much smaller extent than in the mouse (Liska et al., 2022). By contrast, in area 8A of the prefrontal cortex (PFC) of a macaque performing a cognitive task, uninstructed movements clearly modulated, but did not dominate, neuronal activity, although cognitive signals were reliable on the single-trial basis, regardless of whether the uninstructed movements were large or small (Tremblay et al., 2022). These studies raise the possibility that, in primates, the degree of influence that movement-related signals and cognitive signals have on neuronal activity may vary widely across cortical areas, presumably in accordance with their standing in the hierarchy of information processing, and/or depending on their corresponding sensory modalities and somatotopies. Although the somatotopy of the motor and somatosensory cortex is known in both mice and non-human primates (Burish et al., 2008; Burman et al., 2008; Hira et al., 2015; Krubitzer and Kaas, 1990), it is unclear to what extent orofacial and limb movements contribute to activity in these areas in non-human primates. In addition, it has not been examined in primates how the value of an expected reward (a primitive cognitive signal that should largely affect the following operant response) is represented in widespread cortical areas after excluding neuronal activity associated with task-irrelevant movements and operant behaviors (Roesch and Olson, 2003).

In a previous study, in which we conducted calcium imaging in mice during a classical conditioning task with two auditory cues assigned to different reward probabilities, we found that the anticipatory lick dominated the activity throughout the dorsal neocortex after the auditory cue presentation (Kondo and Matsuzaki, 2021). Nevertheless, when imaging cortical areas, reward-predicting cues are most strongly represented in the secondary motor cortex (M2), which is thought to correspond to the premotor cortex (PM) in primates. In this study, we applied the same classical conditioning task to the common marmoset, and conducted unprecedented wide-field one-photon calcium imaging of PM, primary motor cortex (M1), primary and secondary somatosensory cortex (S1 and S2), posterior parietal cortex, and primary auditory cortex (A1). We address: 1) the extent to which orofacial and limb movements modulate cue-triggered activity in these areas of the marmoset, and 2) whether and how the values of sound cues are represented in marmoset PM.

## Results

### Head-fixed marmosets show anticipatory licks during a classical conditioning task

We trained two adult marmosets (Marmoset 1 and Marmoset 2) to perform a classical conditioning task under head-fixation (Fig. 1a). Two sinusoidal tones were associated with different probabilities of reward delivery: Tone A (16 kHz) was associated with a high (70%) reward probability (cue A trials), whereas Tone B (6 kHz) was associated with a lower (30%) reward probability (cue B trials) (Fig. 1b). In each trial, either of the tones was randomly presented for 2 s (cue period), followed by a 1-s delay period, then the reward was delivered according to the reward probability associated with the tone. Movements of the marmosets’ eyes, body, and face were monitored with three cameras (Fig. 1a).

**Figure 1:**
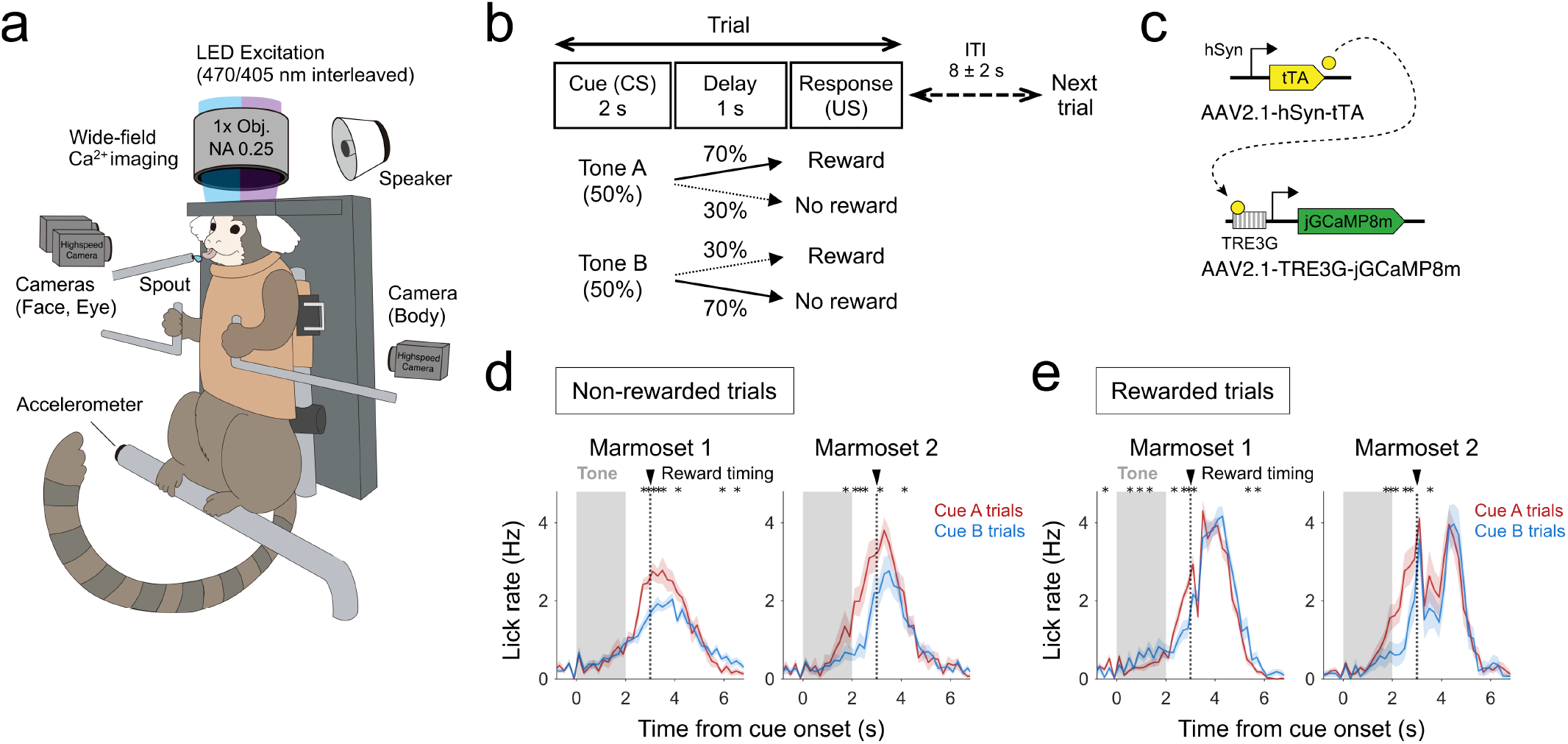
Wide-field calcium imaging of behaving marmosets performing a classical conditioning task. **a** Overview of the experimental setup. Each marmoset was head-fixed and wore a jacket as single-photon wide-field calcium imaging was performed. A speaker behind the animal generated auditory stimuli. A lick spout was prepared in front of the animal to deliver the water reward. Three high-speed cameras were used to monitor the behavioral activity of the animal (eye camera, face camera, and body camera). An accelerometer was attached to the footrest of the animal, to act as an electric sensor of foot motion. **b** Schematics of the task. The head-fixed animal was conditioned to two types of tones with different reward probabilities. ITI, inter-trial interval. **c** A schematic of calcium sensor expression. A tTA-based amplification system was used to express jGCaMP8m. **d**,**e** Conditioned lick behavior during non-rewarded (**d**) and rewarded (**e**) trials. Lick events were recorded using a lick sensor attached to the lick spout. Trial-aligned, 200 ms-binned lick traces of the animals used in the study are shown. Note the existence of anticipatory licks around the time of the reward delivery. Marmoset 1, *n* = 10 sessions; Marmoset 2, *n* = 9 sessions. The mean ± SEM of session-average traces is shown. \**P* < 0.05, Wilcoxon signed-rank test between cue A and cue B trials.

After 52 sessions for Marmoset 1 and 37 sessions for Mar-moset 2, we conducted a craniotomy over the frontoparietal and superior temporal cortices, injected adeno-associated viruses (AAVs) carrying a tetracycline-inducible jGCaMP8m expression system (Ebina et al., 2018; Kimura et al., 2023; Obara et al., 2023; Sadakane et al., 2015; Zhang et al., 2023) (Fig. 1c), and set two glass windows: the first one (14.2 × 7.2 mm) over the frontoparietal cortex and the second one (8.2 × 4.2 mm) over the superior temporal cortex including the lateral sulcus (Supplementary Fig. 1a–c,e). Prior to virus injection, we conducted intracranial micro-stimulation (ICMS) to determine the border between M1 and the caudal part of the dorsal PM (PMdc or 6DC) (Supplementary Fig. 1c,e) (Ebina et al., 2024, 2019). The location of each dorsal neocortical area was inferred according to the location of this border and the marmoset atlas (Paxinos et al., 2012). Estimation of auditory areas took place after we confirmed GCaMP expression; we examined calcium responses through the second glass window while we presented sinusoidal tones of 1–24 kHz. The location of A1 was estimated on the basis of the resulting tonotopy map and the location of the lateral sulcus (Aitkin et al., 1986; Obara et al., 2023; Tani et al., 2018) (Supplementary Fig. 1d,f).

After expression of GCaMP reached a sufficiently high level, we conducted one-photon imaging through the two glass windows (Supplementary Fig. 1b) in 10 sessions (one per day) for Marmoset 1 and 9 sessions (one per day) for Marmoset 2. In these imaging sessions, anticipatory licking gradually increased during the cue period and further increased during the delay period in both animals (Fig. 1d). The lick rate returned to the baseline level around 6 s after the onset of the tone presentation. The anticipatory lick rates around the reward timing were higher in cue A trials than in cue B trials (Fig. 1d). These results imply that the animals successfully learned this classical conditioning task: they expected the reward after the auditory cue presentation, and recognized the difference in the reward probabilities associated with the two cues.

### Calcium responses of the frontoparietal and primary auditory cortices during the classical conditioning task

During the imaging sessions, GCaMP fluorescence was simultaneously recorded from the two glass windows (Fig. 2a,b). We focused on the changes in the cortical activity after the tone presentation (Fig. 2c,d and Supplementary Fig. 2a,b). Several prominent area-specific features were shared in common between the two animals: (1) a large increase or a large decrease in A1 activity was observed during tone presentation, depending on the type of the tone; (2) the activity of the medial part of the rostral part of the dorsal PM (PMdr or 6DR) gradually increased from the onset of the tone presentation until around the timing of the reward delivery, and then decreased; (3) a reduction in activity was observed in the central and lateral parts of PMdr, beginning at around the time of reward delivery and lasting for 2–3 seconds.

**Figure 2:**
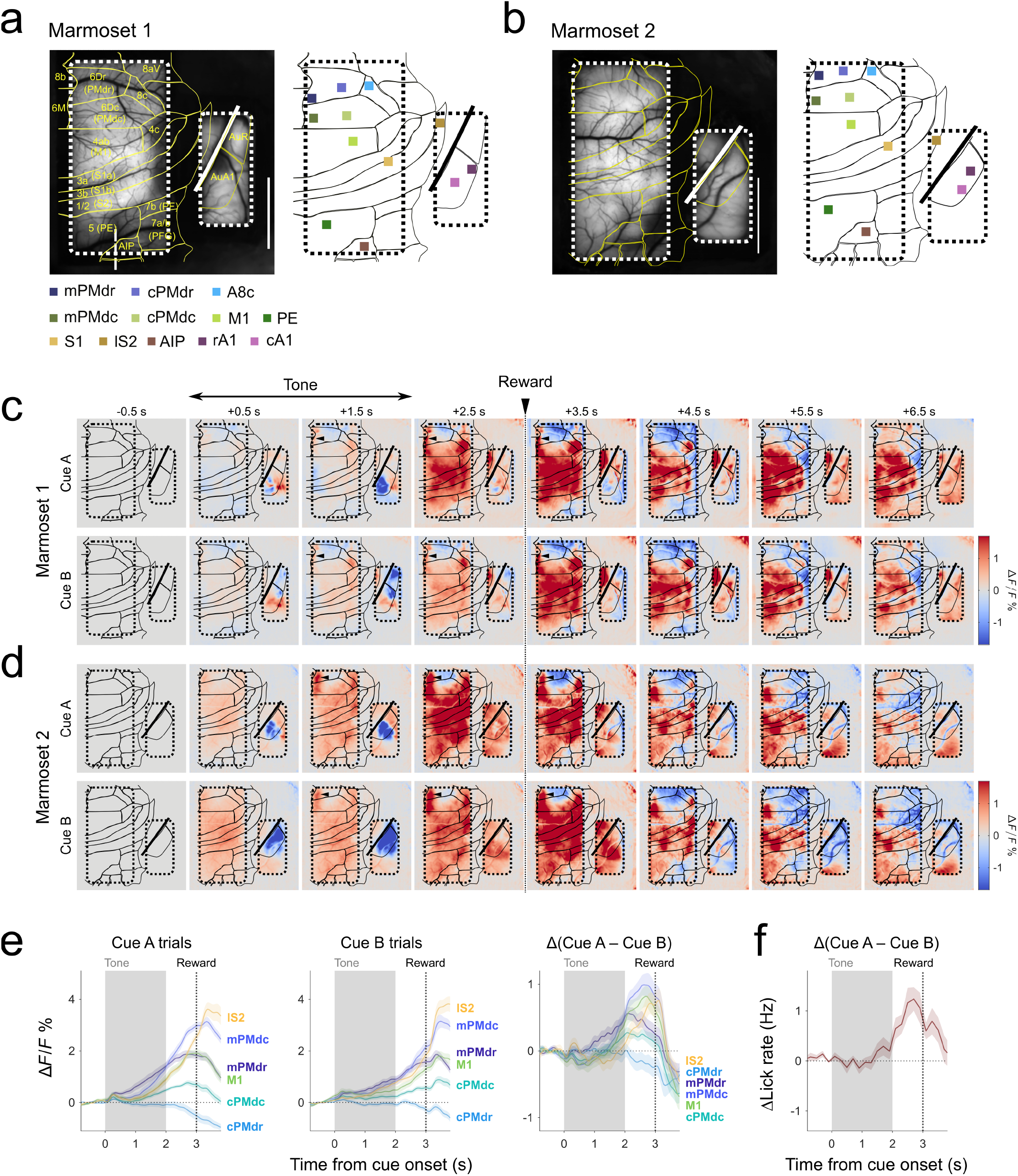
Neocortical calcium responses to the two conditioned stimuli. **a**,**b** Expression of jGCaMP8m in the imaging fields of Marmoset 1 (**a**) and Marmoset 2 (**b**). The mean fluorescence image over the imaging sessions is shown on the left for each animal. The right panels indicate the selection of arbitrary ROIs. The ROIs were selected so that their positions matched between the two animals. For PMdr and PMdc, ROIs were set on the caudal side of each area. Marmoset 1, *n* = 10 sessions; Marmoset 2, *n* = 9 sessions. Scale bars, 5 mm. **c**,**d** Trial-averaged calcium responses in cue A (top) and cue B (bottom) trials in Marmoset 1 (**c**) and Marmoset 2 (**d**). The baseline-subtracted responses of individual trials were weighted so as to balance the unequal numbers of rewarded and non-rewarded trials. Arrowheads on panels indicate the medial region of the PMdr (mPMdr). Marmoset 1, *n* = 10 sessions; Marmoset 2, *n* = 9 sessions. **e** The calcium responses of ROIs to the two conditioned tones. *n* = 19 sessions from two animals were pooled and are shown. ROIs were taken from areas that responded to tones before the timing of reward delivery. The responses to cue A (left), cue B (center), and the differences between the two (right), are shown. Each line represents the mean baseline-subtracted response of the corresponding ROI, indicated as the label bearing the same color. Shaded areas around the traces represent the SEM of session-mean responses. The imbalance in the number of trials is corrected in the same manner as in (**c**) and (**d**). **f** Differences in anticipatory lick rates during cue A and cue B trials, shown in a similar manner as in the right-most plot in **e**. *n* = 19 sessions from two animals were pooled and are shown.

To further examine the differences in activity changes across multiple cortical areas, we set 12 regions of interest (ROIs) as representative areas, as shown in Fig. 2a,b. Two ROIs were set for PMdr and two for PMdc because of the qualitatively different response patterns between the medial and central parts of these areas: the medial part of PMdr (mPMdr) and the medial part of PMdc (mPMdc) showed prominent increased activity during the cue and delay periods, in contrast to the central part of PMdr (cPMdr) and the central part of PMdc (cPMdc). We also set rostral and caudal ROIs in A1 (rA1 and cA1) because these subregions showed opposite tone selectivity during the cue period (Fig. 2c,d and Supplementary Fig. 2a,b). The ROI for S2 was set at its most lateral part (lS2) to capture responses related to tongue movement (Burish et al., 2008; Krubitzer and Kaas, 1990; Nomura et al., 2024; Schaeffer et al., 2019b). For the other areas, one ROI per area was assigned (area 8c [A8c], M1, S1, PE, and the anterior intraparietal cortex [AIP]) (Fig. 2a,b).

To examine the potential roles of these ROIs in the representation of reward expectancy, we further narrowed down our focus to their responses during the cue and delay periods. All the ROIs, except for those in rA1, cA1, cPMdr, and A8c, showed increases in activity during the cue and delay periods, with the rise of this increased activity showing varying onsets and amplitudes (Fig. 2c,d and Supplementary Fig. 2a,b). By contrast, cPMdr and A8c showed decreased activity around the delay period. Above all, mPMdr, mPMdc, PMdc, and S1 showed larger increases in activity in cue A trials than in cue B trials at some timepoints during the cue and delay periods (Fig. 2c,d and Supplementary Fig. 2a,b). In Marmoset 1, M1 and PE also had some timepoints during the delay period that showed higher levels of activity in cue A trials than in cue B trials. These results raised the possibility that the cue-selective responses in these ROIs may represent the animal’s expectation of reward delivery. However, when we plotted the time course of the difference in activity between cue A and cue B trials, the resulting traces were reminiscent of the time course of the difference in the anticipatory lick rates (Fig. 2e,f). It thus remained unclear whether these areas represented the reward-predicting cues or lick-related sensory and/or motor signals.

### Changes in orofacial and limb movements during the classical conditioning task

In addition to the anticipatory lick behavior, it seemed possible that movements of other body parts, including eye, limbs, and face, may underlie the neocortical responses during the cue and delay periods. To determine whether other movements occurred after the onset of the cue presentation, we extracted eye, hand, and facial movements from the three videos, and extracted foot movement using an accelerometer (Fig. 3a–f and Supplementary Fig. 3). Facial movements were extracted as the deviation of 18 key-points on the face from their mean positions, and were represented in terms of their principal components (PCs). The time courses of trial-averaged movements indicated that many dimensions of the animal’s behavior were phase-locked to the task events. For example, slight and transient increases in pupil diameter were observed after the cue onset, and these appeared to be common to both marmosets (Fig. 3d). However, many task-aligned tone-selective aspects of their spontaneous behavior, e.g., eye position for Marmoset 1 and hand speed for Marmoset 2 during the cue and delay periods, were specific to individual animals. The existence of these animal-specific task-aligned behaviors needed to be accounted for to reveal the property of the cue-triggered neocortical activity common across the two animals.

**Figure 3:**
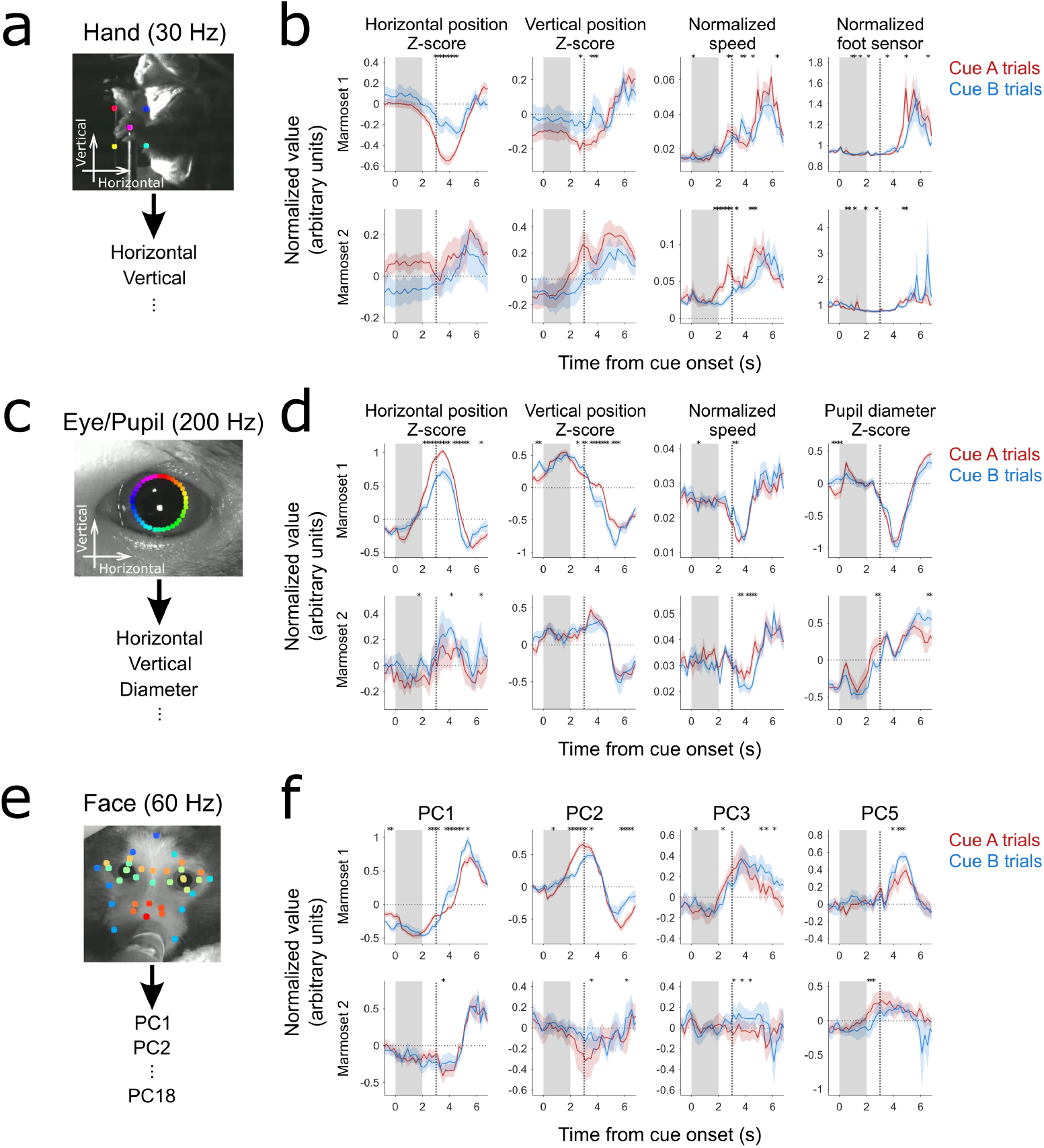
The behavior of the marmosets reflected the task structure. **a**,**c**,**e** Representative images of hand (**a**), eye (**c**), and face (**e**) from the videos, tracked at 30 Hz, 200 Hz, and 60 Hz, respectively. Colored dots represent the key-points being used to extract the position and speed of each part. Face key-points were detected from each frame, and their deviations from the means were decomposed into a set of principal component scores (PC1–PC18). **b**,**d**,**f** Non-baseline-subtracted traces of four hand/foot-related features (**b**), four eye/pupil-related features (**d**), and four face features (**f**) in Marmoset 1 (top) and Marmoset 2 (bottom). Trial-averaged, 200 ms-binned traces aligned to the cue onset (cue A, red; cue B, blue) were session-averaged. The imbalance in the number of trials is corrected. The accelerometer attached to the footrest (not shown) was used as the foot sensor (furthest right in **b**). \**P* < 0.05, Wilcoxon signed-rank test between cue A and cue B trials. Marmoset 1, *n* = 10 sessions; Marmoset 2, *n* = 9 sessions.

### The contributions of movements and the reward-predicting cue to cortical activity

To clarify the contributions of licking and hand, foot, eye, and facial movements and task events (i.e., cue presentation and reward delivery) to the activity of each cortical area, we employed a ridge regression model (see Methods for details). In our design matrix, the input variables (regressors) to the model consisted of the time series of flags for cues (cue A or cue B trials) and the flags for being rewarded, and all the motion-related parameters from our behavioral tracking (Fig. 4a). Time-shifted versions of motion-related parameters were also included in the design matrix to account for the preparatory and feedback activity related to movements. All these different parameters were used to predict the entire time course of the calcium response for each imaging pixel (Fig. 4a and Supplementary Fig. 4a). A ten-fold cross-validation model was used, and the cross-validation-explained variance (cvR^*2*^) was computed to report the performance of each model (Kondo and Matsuzaki, 2021; Musall et al., 2019). On average, 20%–60% of the total fluctuations in all the time series of activity in each of the 12 ROIs were explained by this linear encoding model (Fig. 4b,c and Supplementary Fig. 4a).

**Figure 4:**
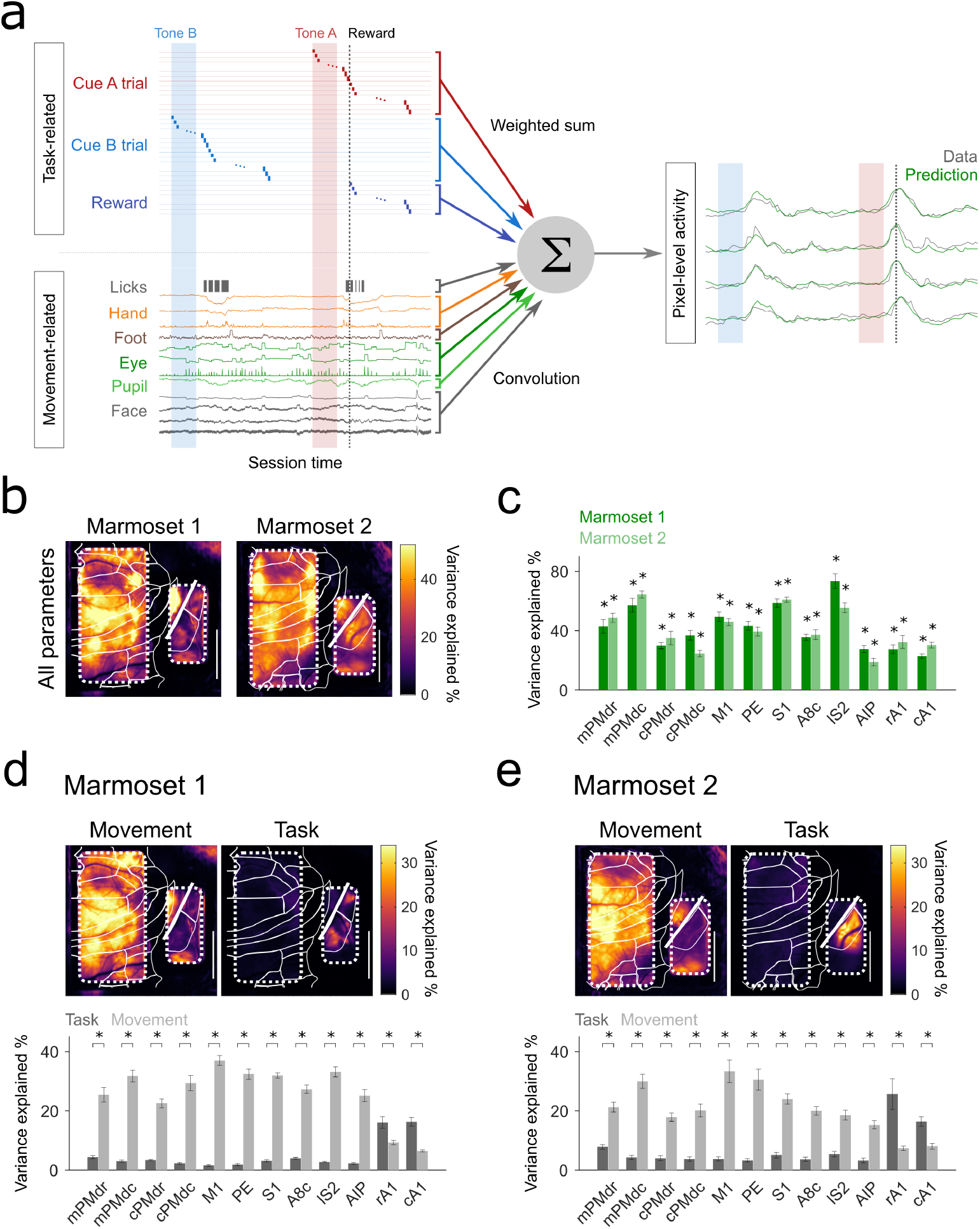
Separation of task- and movement-related neural activity. **a** Schematics of the encoding model. The time series of each pixel was fitted to a linear product of the task-related and movement-related variables (regressors). Task-related flags represent the time periods when the cue A- or cue B-trials were on, and where there was a reward delivery. Movement-related variables were time-shifted to represent 1-s preparatory and 1-s feedback periods. **b** Fraction of the time-to-time variance explained by all the parameters used in the linear encoding models for Marmoset 1 (left) and Marmoset 2 (right). The cvR^2^ values were computed from the time-to-time prediction of calcium signals. The difference in fractions between the full and null models is color-coded in a pixel-by-pixel manner. **c** Comparisons of the ROI-based time-to-time variance explained by all the variables used in the encoding models. Each value was compared against the chance level (\**P* < 0.05, resample test). Marmoset 1, *n* = 10 sessions; Marmoset 2, *n* = 9 sessions. **d**,**e** Fraction of the variance uniquely explained by the task-related and movement-related variables for Marmoset 1 (**d**) and Marmoset 2 (**e**). The unique contribution was defined as the reduction in cvR^2^ values (ΔcvR^2^) after shuffling the columns corresponding to the set of variables under consideration. The top panels represent the comparisons of unique contributions in a pixel-by-pixel manner. The bottom panels compare the task- and movement-related unique contributions in the 12 representative ROIs. Note the unique contributions of the movement-related variables are larger than those of the task-related variables in all the ROIs under consideration. \**P* < 0.05, resample test. Marmoset 1, *n* = 10 sessions; Marmoset 2, *n* = 9 sessions.

We first examined the unique contributions of the distinct sets of variables used in the model (Supplementary Fig. 4b). These unique contributions were computed by calculating the reduction in the explained variance (ΔcvR^*2*^) after time-shuffling the set of variables to be examined. As expected from their calcium responses, A1 areas showed large cue-related ΔcvR^*2*^ values (Supplementary Fig. 4b). Large hand-related unique contributions were detected mainly in M1, S1, S2, and PE (Supplementary Fig. 4b). A large lick-related unique contribution was noted in the lateral side of area 8c, S1, and S2, presumably reflecting the orofacial somatotopy of these regions (Supplementary Fig. 4b). Eye-related unique contributions were prominent around area 8c in Marmoset 1 and a part of the posterior parietal cortical areas in both animals (Supplementary Fig. 4b), presumably reflecting the activity related to saccadic eye motion in these areas (Schaeffer et al., 2019b; Selvanayagam et al., 2019). A slight foot-related unique contribution was observed in the medial side of M1, S1, and PE. The spatial patterns of representation of these movements are in line with the somatotopy of the motor and sensory cortices in non-human primates. By contrast, the cue- and reward-related unique contributions were overall low in the cortical areas, except for A1. Taken together, during the simple classical conditioning with the sound cues, the unique contribution of the total behavioral variables was much larger than that of the total task event variables, except in the auditory cortex (Fig. 4d,e).

### Tones and the difference between the reward expectancies are represented in mPMdr

So far, our analysis of the variance explained by the encoding models has been based on the signals from the full-duration imaging sessions. Although it helps to provide a more accurate estimation of the behavioral contributions, this approach may have underestimated the contribution of task-related events that appeared at a very limited number of timepoints in each imaging session. We therefore examined the time courses of the representations of cue A- and cue B-related variables using 1-s time bins aligned with the cue onset (Fig. 5a). In both marmosets, spot-like patterns of large unique contributions were observed in the medial part of the dorsal premotor areas during the cue and delay periods of the trial (Fig. 5a, arrowheads). More diffuse patterns were detected over a large fraction of somato-motor areas after the reward timing. We then examined the amount of information related to reward-predicting cues by computing the unique contribution of cue-related variables in our representative ROIs during the cue and delay periods (0–3-s period from the cue onset). Although the cue-related unique contribution was detected in many areas, it was significantly larger in mPMdr than in the other areas, except for A1 (Fig. 5b). We also compared the differences between cue A- and cue B-related unique contribu-tions as the ΔVariance (Cue A – Cue B) (Fig. 5c,d). The difference in unique contributions during the cue and delay periods was larger in mPMdr than in the other areas except for A1, although this difference was not found to be significant (Fig. 5d). Taken together, these results suggest that even when we accounted for the contribution from the marmoset’s movements, mPMdr held a significant fraction of information related to reward-predicting cues.

**Figure 5:**
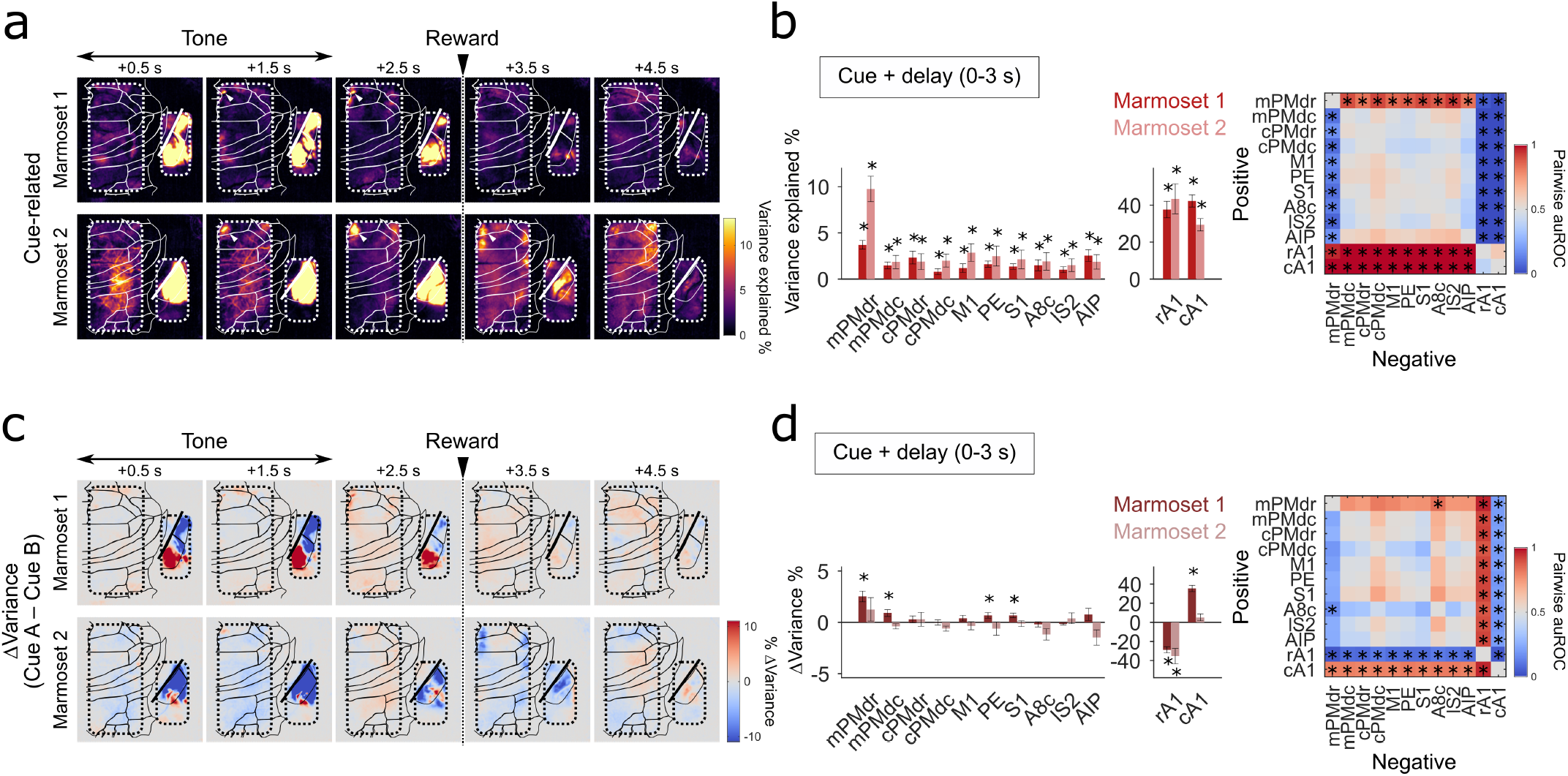
Cue-related modulation of mPMdr before the reward timing. **a** The unique contribution of the cue-related variables was computed separately for individual time bins along the progression of the trial. The pixel-by-pixel means of ΔcvR^2^ values over the sessions are color-coded and shown as images. Arrowheads on panels indicate the medial region in the PMdr (mPMdr). **b** Left, the unique contributions of the cue variables during the cue and delay periods for each ROI. Right, matrix of pairwise area under the receiver operating characteristics curve (auROC) values. \**P* < 0.05, resample test (left; comparison with the chance levels), and resample test with Holm-Bonferroni correction (right). **c** Similar to **a** but for the differences between cue A-related and cue B-related unique contributions, ΔVariance(Cue A – Cue B), along the procession of the trial. **d** Quantification of ΔVariance(Cue A – Cue B) for the representative ROIs during the cue and delay periods of the trial. The values and comparisons are shown in the same manner as in **b**. \**P* < 0.05, resample test (bar plots), and resample test with Holm-Bonferroni correction (matrices). Marmoset 1, *n* = 10 sessions; Marmoset 2, *n* = 9 sessions.

We further addressed the possibility that the cue-related unique contribution in mPMdr may be explained solely in terms of pure sensory responses. To this end, we performed a separate set of auditory stimulation experiments, in which we presented the animals with task-relevant (16 and 6 kHz) and task-irrelevant (1, 2, 4, 8, 12, and 24 kHz) tones (Supplementary Fig. 5). All the tones were presented without any association with reward delivery in these experiments. The responses of rA1 and cA1 ROIs to tone presentation resulted in similar patterns of activity after tone offset, although the amplitude of responses during tone presentation in cA1 was smaller for cue tones than for non-cue tones (Supplementary Fig. 5a, right panel), which may have been related to the animal’s engagement with task-related tones (Otazu et al., 2009). By contrast, cue tones appeared to elicit larger responses in mPMdr than non-cue tones did (Supplementary Fig. 5a, left panel). To remove the effect of movements from these responses, we used linear encoding models in a similar manner as for the classical conditioning task. Quantification based on the unique contributions during the 3-s period after tone onset confirmed our results of stimulus onset-aligned calcium responses: task-related tones were overrepresented in mPMdr in comparison with task-unrelated tones, whereas responses to task-unrelated tones tended to be larger in A1 (Supplementary Fig. 5b). These results are consistent with our hypothesis that mPMdr maintains task-relevant representations rather than purely sensory ones.

It should also be noted that when we examined the cue-related unique contributions during the response period (3-s period from the reward timing, or 3–6 s from the cue onset), the prominent representation of cue information in mPMdr disappeared (Fig. 5a and Supplementary Fig. 6a,b). This implies that the role of mPMdr in maintaining the representation of reward-predicting cues is primarily restricted to the time period before the reward timing.

### Somatotopic representation of orofacial and limb movements

To further assess the effects of movements on cortical activity, we examined the extent to which the movement-related variables explained the activity of ROIs during the cue and delay periods (Fig. 6a–f). Although almost all ROIs significantly represented individual movements, the individual magnitudes of representation differed across ROIs. Licking was strongly represented in S1 and lS2, but was weakly represented in AIP, rA1, and cA1 (Fig. 6a). Hand movement was strongly represented in cPMdc, M1, and PE (Fig. 6b). Foot movement was weakly represented across the ROIs being examined (Fig. 6c). Eye movement was well represented in mPMdr, cPMdr, A8c, and AIP (Fig. 6d). Pupil diameter and face movement were relatively broadly represented (Fig. 6e,f). When we examined the extent to which the variables explained the neuronal activity during the response period, the results were very similar (Supplementary Fig. 6c–h). These results were also qualitatively consistent with the results of the maps of the contribution of each movement for the entire time series (Supplementary Fig. 4b). These results suggest that although some movements were phase-locked to the cue presentation, the extent to which these movements uniquely affected the neuronal activity was not specific to any phases of the trial (i.e., the cue, delay, or response periods).

**Figure 6:**
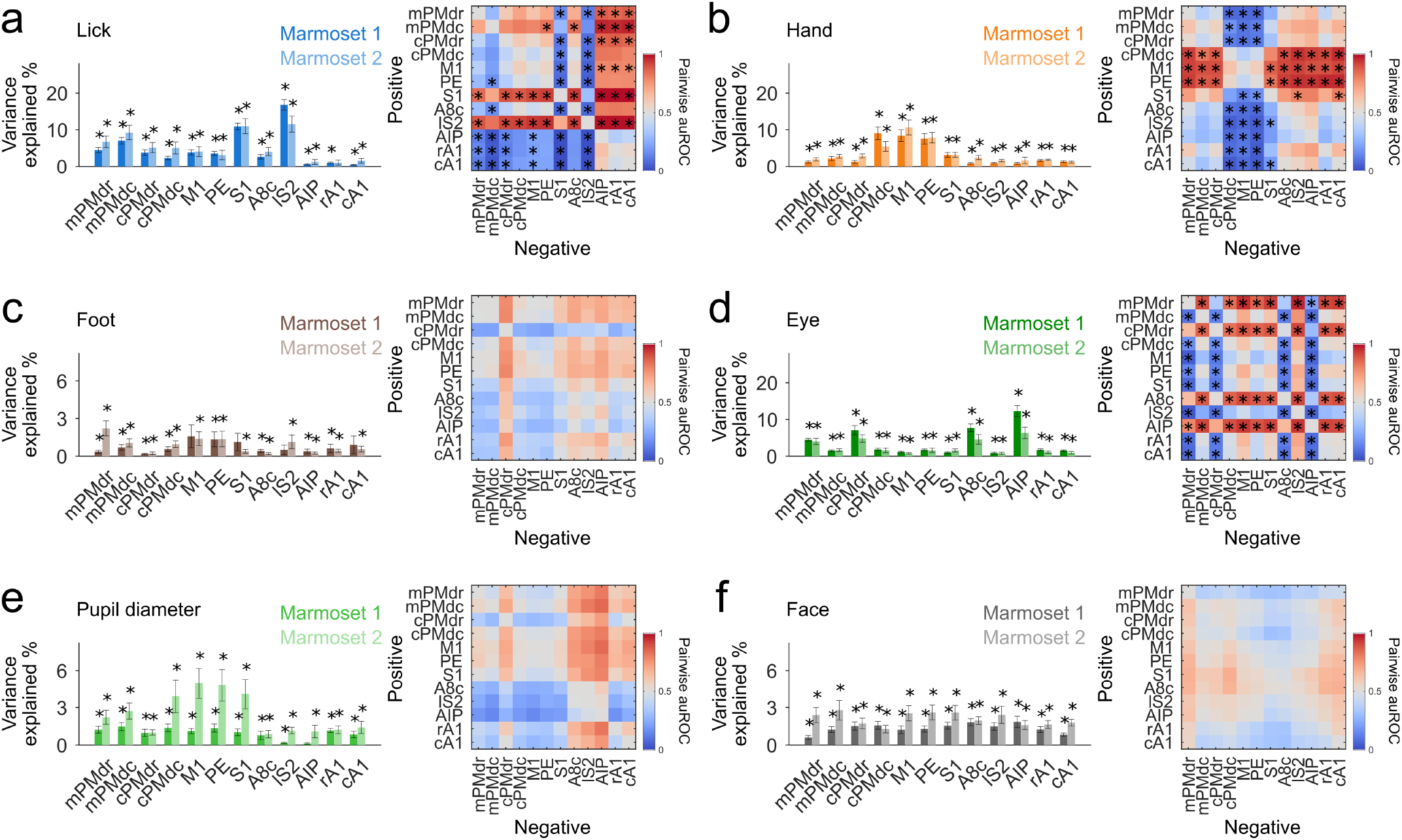
Unique contributions of movement-related variables during the cue and delay periods. **a-f** Left, unique contributions of the six movement-related variables: lick (**a**), hand (**b**), foot (**c**), eye (**d**), pupil diameter (**e**) and face (**f**) during the cue and delay periods for each ROI. Marmoset 1, *n* = 10 sessions; Marmoset 2, *n* = 9 sessions. Right, matrix of pairwise auROC values. Data from *n* = 19 sessions from two animals were pooled and are shown. The values were color-coded based on the corresponding auROC values. \**P* < 0.05, resample test (left), and resample test with Holm-Bonferroni correction (right).

We also examined the temporal aspects of movement-related modulation on neuronal activity. Most of the model weights for lick, hand, foot, and eye movements increased before the movement timing and peaked after the movement timing, presumably reflecting the time courses of their corresponding motor-and sensory-related signals (Supplementary Fig. 7a–d). The lick-related weights in S1 and lS2 showed large peak values (Supplementary Fig. 7a), consistent with the robust lick-related representations in these areas. By contrast, some other areas showed negative weights. In particular, cPMdr and A8c, which showed decreased activity during the delay period of the trial, showed negative weights even before the lick onset. This result suggests that suppression of activity occurs in these areas from some time before any detected lick activity, potentially being related to preparation or intention to perform licks. Interestingly, the time course of the weight for the pupil diameter was similar across all areas, and it had a characteristic transient peak that occurred before the time of change in pupil diameter (Supplementary Fig. 7e). This suggests that the manner in which the pupil diameter is related to the dorsal cortical activity is different from the way the other movements contribute to it.

### mPMdr and cPMdr are involved in distinct functional networks in the dorsal cortex

Our results revealed distinct representations of task- and movement-related information in mPMdr and cPMdr (Figs 5, 6; Supplementary Fig. 7). To examine whether these distinct representations were related to different functional cortical networks, we computed seed-based correlations (Nomura et al., 2024). The subnetwork with the cPMdr seed showed a different pattern from the subnetworks seeded with the other ROIs in PM (Fig. 7a,b). The pattern of the latter subnetworks was similar and extended through M1, S1, and S2 to the rostral part of PE. To examine the correlational structures of the 12 ROIs altogether, we performed hierarchical clustering using correlation-based metrics (Fig. 7c–e). The dendrogram based on the activity during the cue and delay periods showed three major clusters (Fig. 7c): one consisting of M1, mPMdc, cPMdc, S1, PE, and mPMdr, another consisting of cPMdr, area 8c, and the AIP, and the third one consisting of rA1 and cA1. We also built dendrograms based on non-trial activity (Fig. 7d) or movement-related encoding model weights (Fig. 7e), and found that their structures were very similar to the one based on trial-related activity. These results suggest that these functional connectivity patterns reflect the somatotopic modular organization of the neocortex. Our results also suggest that mPMdr and cPMdr participated in different sets of functional subnetworks, without the distinction of the within-trial or out-of-trial contexts of classical conditioning.

**Figure 7:**
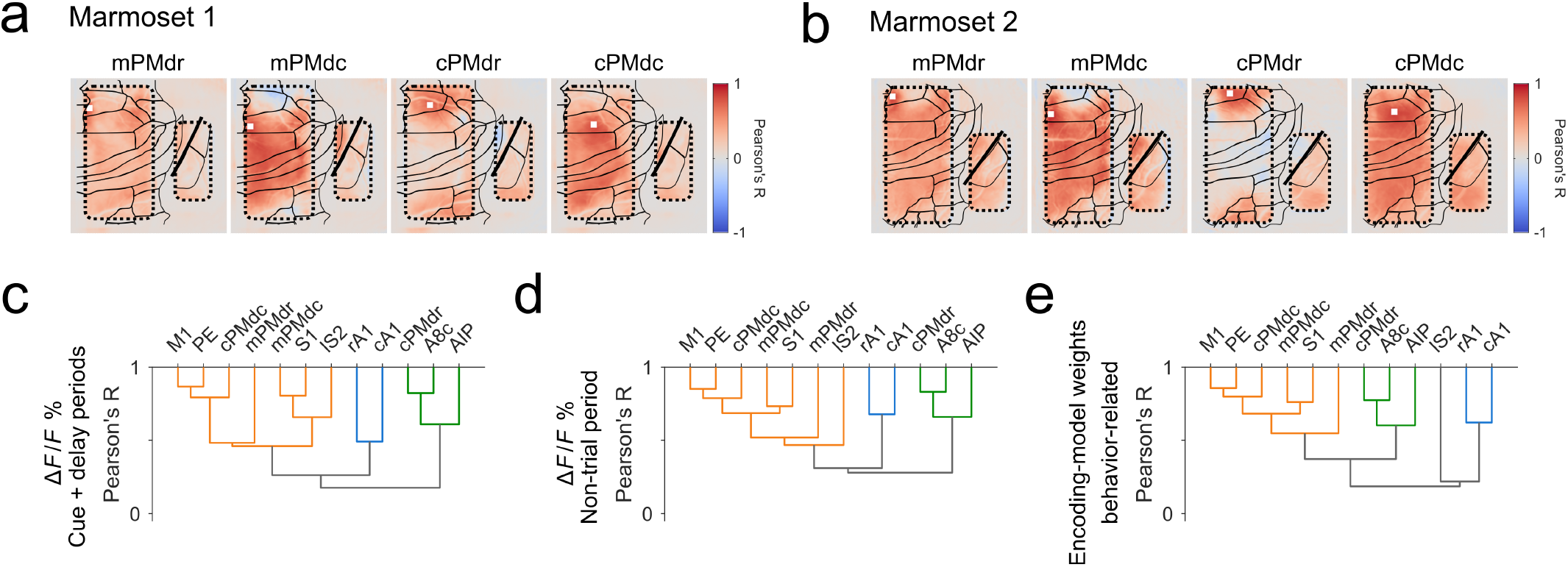
Unique contributions of movement-related variables during the cue and delay periods. **a**,**b** Session-averaged correlation maps with the seed of each of the four ROIs in PMdr and PMdc in Marmoset 1 (**a**) and Marmoset 2 (**b**). The label to each panel and the location of the white rectangle correspond to the ROI being used as the seed. A linear regression was performed for each pixel using the full imaging session time courses. The Pearson’s correlation coefficient, being averaged over the imaging sessions, is shown in a pixel-by-pixel manner as a color-coded image. Marmoset 1, *n* = 10 sessions; Marmoset 2, *n* = 9 sessions. **c** A dendrogram of the ROIs was computed and drawn based on the temporal correlation during the cue and delay period of the trial. Data from *n* = 19 sessions from two animals were pooled and are shown. The cluster was colored when Pearson’s correlation coefficient was above the arbitrary threshold of 0.4. The orange subset of branches corresponds to the subnetwork containing M1 and S1. The green subset corresponds to the subnetwork containing cPMdc and AIP. The blue subset corresponds to the subnetwork containing rA1 and cA1. **d** A dendrogram was computed, in the same manner as in **c**, and drawn based on the temporal correlation during the non-trial periods in the imaging sessions. **e** A dendrogram was computed, in the same manner as in **c**, and drawn based on the similarity of the movement-related weights of the linear encoding models shown in Supplementary Fig. 7.unique contributions of the six movement-related variables: lick (**a**), hand (**b**), foot (**c**), eye (**d**), pupil diameter (**e**) and face (**f**) during the cue and delay periods for each ROI. Marmoset 1, *n* = 10 sessions; Marmoset 2, *n* = 9 sessions. Right, matrix of pairwise auROC values. Data from *n* = 19 sessions from two animals were pooled and are shown. The values were color-coded based on the corresponding auROC values. \**P* < 0.05, resample test (left), and resample test with Holm-Bonferroni correction (right).

## Discussion

In the current study, we conducted wide-field one-photon calcium imaging of the marmoset dorsal neocortex to reveal how orofacial and limb movements (licking, and hand, foot, eye, and face movements) modulated cortical activity during the presentation of auditory cues that were differently associated with reward delivery. The extent of the modulation varied depending on the cortical areas and body parts involved in the movement. As expected, the frontoparietal cortex responded strongly to each corresponding movement: lateral sensory areas strongly responded to licking; cPMdc, M1, and PE to hand movement; the medial part of M1 and S1 to foot movement; and cPMdr, A8c, and AIP to eye movements. A1 responded very strongly to the auditory cues, but the response to licking and eye movement was very weak, although not absent. A striking difference to the results obtained from mice was that there were regions that showed decreased activity in response to licking: cPMdr and A8c. The reward-predicting cues in our classical conditioning task were strongly represented in a part of PMdr, mPMdr, particularly during the cue and delay periods.

In all previous studies that compared representations between cognitive (or task-related) variables and movements, the cognitive tasks required goal-directed movements to achieve a rewarded state. By contrast, we here used a classical conditioning scheme to minimize the cortical activity and movements that may be related to decision making, planning, and preparation of goal-directed movement. In classical conditioning, anticipatory licking is generally considered a reliable demonstration that the animal recognizes the expected reward. In fact, the two marmosets showed clear anticipatory licking, and the frequency of this anticipatory licking was higher in response to cue A than to cue B. Although both animals exhibited some accompanying movements in response to cue presentations, the time course and extent of these movements was not qualitatively common between the animals. Combining these findings with the results of our auditory stimulation experiments, where task-relevant tones were found to be overrepresented in certain cortical areas compared with non-task-relevant tones, it seems reasonable to consider that the unique contribution of the cue-related variables of our encoding models represented a primitive cognitive signal related to the cue value, but not to the decision making, planning, or preparation of voluntary movements.

### Motor representation in the marmoset dorsal neocortex

Our wide-field imaging results, from the PMdr and area 8c at the anterior limit, to the auditory and posterior parietal areas caudo-laterally, seem to help in unifying the picture of how movements affect the neocortical activity of non-human primates. Our finding that non-task-relevant movements were widely represented in PMdr and area 8c is consistent with a macaque study that showed task-irrelevant movement to be strongly represented in area 8a, which is located anterior to PMdr, although the cognitive signals are also apparently represented in area 8a (Tremblay et al., 2022). At the same time, our finding that movements did not dominate A1 during the tone presentation is consistent with studies in macaques and marmosets that show that bodily movements do not strongly affect V1 activity during visual stimulus presentation (Liska et al., 2022; Talluri et al., 2023). Taken together, it is likely that in non-human primates, the effects of movements on neocortical activity depend on the modality of individual areas. By contrast, in our previous study using mice and the same classical conditioning task, licking (and other body movements associated with it) dominated the activity of all dorsal cortical areas, including frontal, auditory, and primary visual areas (Kondo and Matsuzaki, 2021), implying that the modality specificity of movement-dependent modulation is much less prominent in mice. This difference is likely to be related to the ethological differences between mice and marmosets. Mice frequently move their limbs, whiskers, and mouths when they explore their surroundings, whereas by contrast, non-human primates that live in trees may have evolved to process visual and auditory information more precisely and at a higher level, often without moving their limbs or mouths. Even when such non-human primates move their bodies, they may process audiovisual information in their lower sensory cortices, separately from the processing of movement information.

The strength of motor representation in M1, S1, and lS2 corresponded well with the somatotopy that has been revealed by ICMS-based mapping and sensory mapping (Burish et al., 2008; Burman et al., 2008; Ghahremani et al., 2017; Krubitzer and Kaas, 1990; Selvanayagam et al., 2019). At the same time, the time courses of the model weights for pupil diameter were relatively similar across all imaging areas. The transient increase in pupil diameter after the onset of cue presentation may reflect cue-triggered arousal and an increase in attention, which are regulated in part by adrenergic and cholinergic neurons (Joshi and Gold, 2020; Reimer et al., 2014). Considering that these neurons broadly project their axons over the cortex and their cortical activity preceded the pupil dilation (Reimer et al., 2016), it would be reasonable to presume that they modulated pupil diameter and dorsal neocortical activity in different latencies.

### Representation of reward-predicting cues in medial PMdr

Area 6M is located medially to PMdr and PMdc, and its rostral and caudal parts have been shown to anatomically correspond to pre-SMA and SMA, respectively (Bakola et al., 2021). However, neither pre-SMA-like nor SMA-like neuronal activity have been identified in marmosets. In the macaque, neurons in pre-SMA, but not in PMdc, were reported to show sustained activity representing the perceptual signal, to maintain the information from the preceding task-related sensory stimulus (Hernández et al., 2010). Another macaque study showed that in a delayed saccade task, the expected reward is strongly represented in pre-SMA and PMdc (Roesch and Olson, 2003). The human pre-SMA was also shown to be involved in visuo-motor and audio-motor associations (Kurata et al., 2000; Picard and Strick, 2001; Sakai et al., 1999). Our results, showing that marmoset mPMdr retains information about reward-predicting cues until reward delivery, parallel the previously reported roles of pre-SMA in old-world primates: context-dependent association of sensory stimuli, motor outputs, and rewarding outcomes. It is important to note that our classical conditioning task, unlike the aforementioned studies, does not require any goal-directed movements of the forelimb or eyes. Thus, it would be more unlikely that the observed task-related information in mPMdr represents motor-outcome association, rather than stimulus-outcome association. Along with our results from the encoding model analysis, it would be reasonable to propose that the mPMdr of marmosets may play a functional role being homologous to the pre-SMA in other non-human primates, and that among the higher-order motor areas, the mPMdr and/or pre-SMA of marmosets is the site that is the most modulated by reward expectancy. Further studies are necessary to determine the extent to which the representation of the reward-predicting cues extends anteriorly (area 8a) and medially (area 6M) in the marmoset (Burman et al., 2006).

Our functional correlation mapping showed that mPMdr and cPMdr participated in distinct functional subnetworks; the connection between mPMdr and intraparietal areas was weaker than that between cPMdc and intraparietal areas, with mPMdr appearing to be involved in a subnetwork with M1 and S1. This is consistent with anatomy-based reports that 6M neurons innervate M1 more strongly than do PMdr neurons (Burman et al. 2014a), and that PMdc receives the inputs from area 6M more than PMdr does (Burman et al., 2014b). Resting-state functional magnetic resonance imaging (fMRI) also demonstrated that one functional connectivity-defined cluster that mainly included M1 also included the middle parts of PMdr and PMdc, whereas another cluster connecting with the intraparietal areas included area 8c and the lateral parts of PMdr and PMdc (Schaeffer et al., 2019a). Considering that this fMRI experiment was conducted in a lightly anesthetized state, these resting-state functional clusters could be considered to be independent of any movement-related modulations, and therefore presumably reflected intrinsic functional modules in the marmoset neocortex. Being considered alongside these previous reports, our results add to the idea of intrinsic functional modules that retain their basic structure when the animal undergoes cognitive tasks.

In our previous study using the same classical conditioning task in mice, dorsomedial M2 showed the strongest representation of the reward-predicting cue in the dorsal frontal neocortex (Kondo and Matsuzaki, 2021). This suggests that in both rodents and primates, the medial part of the premotor cortex may be the primary site in the dorsal neocortex in which the sensory input-based level of motivation is represented, and that in turn, the medial part of the premotor cortex modulates the planning and execution of the animal’s upcoming behavioral outputs. It remains to be verified whether this area solely processes auditory-related contextual information, or whether it alternatively is a more general site processing context-relevant information, as the SMA and pre-SMA do in old-world primates.

### Reduction in the activity of central PMdr and area 8c during licking

Unexpectedly, we observed a robust reduction in the activity of cPMdr and A8c before and during the response period, presumably when the animal was licking. By comparison, no area in the mouse dorsal cortex showed decreased activity during licking (Kondo and Matsuzaki, 2021; Musall et al., 2019). In accord with these reports in mice, we previously observed a clear increase in the activity of PMdr while headfixed marmosets performed visual cue-triggered forelimb movement (Ebina et al., 2024). PMdr and area 8c connect with intraparietal areas, and they have been considered to specifically process ocular and/or visual information. It is possible that in our current study, the state of the animal, presumably related to licking and/or reward consumption, resulted in suppression of the activity in these areas during the response period of the trial. This possibility is supported by the finding that locomotion negatively modulates activity in marmoset V1 (Liska et al., 2022). The functional relevance of these movement-related modulations of cortical activity, particularly those in the negative direction, remains to be further examined.

### Advantages and limitations of the marmoset research

The field of view (FOV) of the two windows used in the present study was 136.9 mm^*2*^ (14.2 × 7.2 mm + 8.2 × 4.2 mm). To our knowledge, this is the largest FOV to have been used for one-photon calcium imaging of the mammalian brain. The implanted glass windows were inevitably tilted slightly towards each other (∼20 degrees at the largest), and therefore the optical axis of the microscope was not accurately adjusted perpendicular to the cortical surface. However, we believe that this misalignment had little impact on our results because we were able to robustly extract area-specific signals from both windows. We further expect that it would be possible to achieve even larger imaging fields, e.g., fields including the prefrontal cortex or the occipital cortex, whose cortical surface angle differs substantially from that of the frontoparietal cortex, by installing multiple one-photon microscopes with different tilt angles, as has already been demonstrated for two-photon imaging of mouse brain (Wagner et al., 2019). Although the FOV of two-photon imaging is much smaller, two-photon calcium imaging with a FOV of 3 × 3 mm at a frame rate of 7.5 Hz has already been achieved in the marmoset neocortex (Ebina et al., 2024). In combination, application of these cortex-wide imaging techniques to the marmoset neocortex should help us understand cortex-wide inter-areal information processing in non-human primates. Since there is no clear intraparietal sulcus in the marmoset neocortex, different areas across the posterior parietal cortex should be simultaneously imaged. Higher and primary auditory cortices have also been simultaneously imaged (Obara et al., 2023; Song et al., 2022). In future studies, it should be possible to examine the cellular basis of end-to-end information flow in the marmoset neocortex, from the processing of visual or auditory inputs in the primary sensory areas, through the representation of cognitive processes in the cortical association areas, to the motor areas where integrated information is converted into movement outputs, including vocalization and other social interactions of marmosets (Miller and Wang, 2006; Pomberger et al., 2019; Song et al., 2022; Voelkl and Huber, 2000; Zanini et al., 2023).

On the flipside, the less pronounced sulcal structures of the marmoset neocortex imply a drawback, in that cortical areas cannot be distinguished by morphological landmarks that are easily detectable on visual inspection. Failure to take into account animal-to-animal differences in the sizes and positions of individual neocortical areas would lead to erroneous attributions of activity to underlying neocortical areas. To alleviate such issues, careful registration of each brain to the standard reference brain would be necessary, possibly by conducting MRI (Hata et al., 2023). Optical imaging of intrinsic signals would also be effective for accurately identifying hierarchical sensory areas (Song et al., 2022). Furthermore, the repertoires of available motor and cognitive tasks are still limited compared with those available for macaques; it is necessary to develop behavioral paradigms that fit well with the ethology of marmosets (Matsuzaki and Ebina, 2020; Mitchell et al., 2024).

## Acknowledgements

We thank M. Hirokawa and Y. Takahashi for animal handling and S.-I. Terada for making glass window assemblies. This work was supported by Grants-in-Aid for Scientific Research on Innovative Areas (17H06309 to M.M., 19H05307 to T.E., and 21H00302 to T.E.), for Transformative Research Areas (A) (22H05160 to M.M., 22H05157 to K.I., and 23H04977 to T.E.), for Scientific Research (A) (19H01037 and 23H00388 to M.M.), and for Scientific Research (B) (20H03546 to T.E.) from the Ministry of Education, Culture, Sports, Science, and Technology, Japan; AMED (JP19dm0207069 to M.M.; JP15dm0207001 to T.Y. and M.M.; JP18dm0207027 to M.M.; JP19dm0107150 to M.M.; JP19dm0207085 to T.E. and M.M.; JP22dm0207077 to M.T.; JP23wm0625001 to M.M.); and the Tokyo Society of Medical Sciences (to T.E.). This work was also supported by the program for Brain Mapping by Integrated Neurotechnologies for Disease Studies (Brain/MINDS) from AMED under Grant number JP21dm0207111. This paper was typeset with the bioRxiv word template by @Chrelli: www.github.com/chrelli/bioRxiv-word-template.

## Author contributions

K.S., M.K., and M.M. designed the experiments. M.K. mainly conducted the experiments. Y.H. trained the marmosets. M.K. and T.E. conducted the animal surgery. K.S. and M.K. analyzed the data. K.S., M.K., and T.E. set up the task apparatus for the experiments. M.T., A.W., K.I., M.T., and T.Y. designed and prepared rAAVs. K.S., M.K., and M.M. wrote the paper, with comments from all authors.

## Methods

### Animals

All experiments were approved by the Institutional Animal Care and Use Committee of the University of Tokyo. One male (Marmoset 1) and one female (Marmoset 2) laboratory-bred common marmosets (Callithrix jacchus) were used in the present study. Their ages were 25 and 21 months and their weights were approximately 350 g when the habituation started. Both marmosets were kept under a 12:12-hour light-dark cycle and the light phase was started at 8 AM. Neither were used for any other experiments prior to the present study.

### Virus production

The AAV plasmid of the human synapsin I promoter (hSyn)-tetracycline-controlled transactivator 2 (tTA2s) was described previously (Ebina et al., 2019, 2018; Konno and Hirai, 2020). The plasmid for tetracycline response element (TRE)-jGCaMP8m was constructed based on the TRE3G-GCaMP6f vector (Sadakane et al., 2015), by replacing GCaMP6f with jGCaMP8m from pGP-AAV-syn-jGCaMP8m-WPRE (a gift from the GENIE Project, Addgene plasmid 162375) (Zhang et al., 2023). The virus particles, AAV2.1-hSyn-tTA2S, and AAV2.1-TRE3G-jGCaMP8m (Kimura et al., 2023) were produced using the helper-free triple transfection procedure, purified by affinity chromatography (GE Healthcare, Chicago, IL), and concentrated by ultrafiltration (Amicon Ultra-4 10K MWCO; Merck KGaA, Darmstadt, Germany). Viral titer was determined by quantitative PCR using Taq-Man technology (Life Technologies, Carlsbad, CA).

### General handling and training schedule

Before they underwent headplate implant surgery, the marmosets were first habituated to the experimenters, to wearing a jacket, and to water restriction. Training of the classical conditioning task with head-fixation started more than a week after the headplate implant (days after surgery, Marmoset 1, 23 days; Marmoset 2, 19 days). The animals performed the behavioral task 1–5 days per week, during which their body weight was maintained at approximately 90% of their normal level by restriction of food and water. On the off-duty days, the food and water restrictions were weakened to allow them to return to their normal weight. The behavioral sessions were carried out once per day.

Classical conditioning pre-training sessions started by using 100% reward probabilities for both cues. Probabilities were lowered to 70% (cue A) and 30% (cue B) after a robust anticipatory lick behavior appeared across sessions (Marmoset 1, 38 sessions; Marmoset 2; 40 sessions). Craniotomy, intracortical micro-stimulation (ICMS), and virus injection into each animal were conducted after differential anticipatory lick behavior was confirmed for the two tones (Marmoset 1, 52 sessions; Marmoset 2, 37 sessions, after introducing the difference in reward probabilities). Food and water restrictions were lifted several days before these surgical procedures (Marmoset 1, 11 days; Marmoset 2, three days). Classical conditioning training, along with food and water restriction, restarted at least a week after the surgery (Marmoset 1, 11 days; Marmoset 2, 29 days). A set of imaging sessions was started after we confirmed that viral expression was sufficiently high (Marmoset 1, 39 days; Marmoset 2, 38 days, after virus injection).

### General surgical procedures

All the surgeries and viral vector injections were carried out under aseptic conditions described previously (Ebina et al., 2024, 2019, 2018). Each marmoset was placed in a stereotaxic instrument (SR-6C-HT; Narishige, Tokyo, Japan), 0.8%–4.0% isoflurane anesthesia at a flow rate of 1 L/minute was maintained, and the saturation of percutaneous oxygen (SpO2), pulse rate, and rectal temperature were monitored throughout surgery. In the perioperative period, the following medicines were intramuscularly (i.m.) administered: ampicillin (16.7 mg/kg of body weight; Nippon Zenyaku Kogyo, Fukushima, Japan) as an antibiotic, carprofen (Remadile®, 4.4 mg/kg; Zoetis, Parsippany, NJ) as an anti-inflammatory agent, and maropitant (Cerenia®, 1000 mg/kg; Zoetis) as an antiemetic. To avoid dehydration, 10 mL of Ringer’s solution and riboflavin (Bisulase®, 0.02 mL/mL Ringer solution; Toa Eiyo, Fukushima, Japan) were subcutaneously infused.

### Headplate implantation

In the headplate implantation procedure, we depilated and sterilized the scalp, incised it with an external application of lidocaine, and removed the connective tissue to expose the skull. After small screws were anchored to the skull, a headplate was attached to the skull with dual-cured adhesive (Bondmer lightless®, Tokuyama Dental, Tokyo, Japan) and dental resin cement (Estecem II®; Tokuyama Dental, and Superbond®; Sun Medical, Shiga, Japan).

### Craniotomy and durotomy

Craniotomy and durotomy for dorsal and temporal cortices were carried out under anesthesia, and the medications described above were administered with additional intramuscular administration of dexametazone (Decadron®; 0.5 mg/kg; Sandoz pharma, Tokyo, Japan) and subcutaneous administration of 20% D-mannitol (2 g/kg; Yoshindo Inc., Toyama, Japan) to prevent cerebral edema. After the surgery, the exposed cortex was covered with silicone elastomer (Kwik-Sil®; World Precision Instruments, Sarasota, FL) and further covered with dental resin cement (Superbond®).

### Intracortical micro-stimulation (ICMS)

Before rAAV injection into the cerebral cortex, ICMS was conducted in a similar way as has been described previously (Ebina et al., 2024, 2018; Nomura et al., 2024) to identify the boundaries between the primary motor cortex (M1), premotor cortex (PM), and somatosensory cortices. During the ICMS, we anesthetized each marmoset with ketamine (Ketalar®, initial dose 15 mg/kg; additional dose 5 mg/kg; Daiichi Sankyo Propharma, Tokyo, Japan) and xylazine (Seractal®, initial dose 0.75 mg/kg i.m.; additional dose 0.25 mg/kg; Beyer Pharma Japan, Osaka, Japan), and administered atropine (0.050 mg/kg i.m.; FUSO Pharmaceutical Industries, Osaka, Japan) for sialoschesis to prevent respiratory obstruction. A silver reference electrode was immersed in the cerebrospinal fluid on the exposed cerebral surface, then a tungsten microelectrode (impedance 0.5 MΩ, diameter 100 μm; World Precision Instrument) was inserted into the cerebral cortex to a depth of 1.8 mm. Electrical pulses were controlled with an electrical stimulator and isolator (SEN-3301 and SS-203J, Nihon Kohden, Tokyo, Japan). Fifteen 0.2-ms cathodal pulses of 333 Hz were applied. We increased the stimulation currents from 10 to 100 μA in steps of 10 μA until a body-part movement was detected by visual inspection. We determined the rostrocaudal position of the PMdc-M1 border according to the current threshold required for movement generation. Using this border, and information about the distance from the midline, the marmoset cortical map (Paxinos et al., 2012) was overlaid on top of the ICMS map. Alignment of the cortical map with the imaging FOV was performed according to the vasculature patterns of the neocortical surface. It should be noted that we did not take into account the animal-to-animal variability in the size and shape of individual neocortical areas. The area annotation may have therefore been less precise as it got further away from the PMdc-M1 border. For example, a central part of the most posterior region of the imaging glass window, which we inferred to be the AIP, likely contained other intraparietal areas. Thus, the activity of AIP that was related to eye movements probably reflected the activity of the lateral and caudal intraparietal areas that are strongly involved in visual information processing.

### AAV injection and glass window implantation

Mineral oil (Nacalai Tesque, Kyoto, Japan) was backfilled into a Hamilton syringe (25 μL, 702RN, Hamilton Company,Reno, NV) and a quartz pipette with an outer tip diameter of approximately 30 μm (Sutter Instruments, Novato, CA) was prepared. The viral solution containing rAAV2/1-TRE3jGCaMP8m (1.05 × 10^13^ vector genomes [vg]/mL) and rAAV2/1-hSyn-tTA2 (1.75 × 10^13^ vg/mL) was front-loaded with a syringe pump (KDS310; KD Scientific, Holliston, MA). The viral solution was vertically injected into each site at a depth of 500 μm from the cortical surface at a rate of 0.10 μL/minute to a total amount of 0.50 μL. The pipette was maintained in place for 5 minutes before being slowly withdrawn. After injections into multiple sites, the glass-window implant was built with a rectangular glass coverslip (15 × 8 mm and 9 × 5 mm, thickness No. 1, Matsunami glass, Osaka, Japan), and a custom-made outer frame (0.4 mm width, Ebina et al. 2018) was attached to the exposed dorsal and temporal cortices. The inner sizes of the glass windows were 14.2 × 7.2 mm and 8.2 × 4.2 mm for dorsal and temporal cortices, respectively. The thickness of the coverslips was approximately 0.15 mm. The gap between the implant and the cranium was infused with Kwik-sil silicone elastomer and the implant was then fixed with dental resin cement (Super bond).

### Behavioral apparatus and training

The head- and body-restrainer apparatus was as described previously (Ebina et al., 2024, 2018). The marmosets wore a jacket and were seated on the head-fixation frame (O’Hara & Co., Tokyo, Japan). The pitch-and yaw-angles of the frame were adjusted to compensate for the angle mismatch between the surface of the glass windows and the optical perpendicular. The apparatus was placed in a sound-attenuation box (O’Hara & Co.) for training sessions, or a custom-made imaging booth covered with black cloth for imaging experiments, to prevent any unwanted environmental cues. The interior of the sound-attenuation box or imaging booth were illuminated by a white ambient light (40–60 lux) at the ceiling of the box or booth; this helped in restricting the animals’ pupil dilation to observable ranges. Auditory stimuli were delivered from a full-range speaker (FE126NV2, Fostex, Tokyo, Japan) placed behind the head of the animal. To mask environmental sounds, white noise (∼70 dB at the position of the animal’s head) was generated by a custom-made electric circuit, and was constantly presented during all the training and imaging sessions. Reward delivery was controlled with a solenoid valve (HNB1-M5-DC12V, CKD corporation, Aichi, Japan). The animal’s lick behavior was detected by a custom-made lick sensor (Slotnick, 2009) that connected the footrest with the lick spout.

Our classical conditioning paradigm was similar to the one described previously (Kondo and Matsuzaki, 2021): rewards (unconditioned stimulus) were delivered in a probabilistic manner, depending on the tone being presented as the cue (conditioned stimulus). A lick spout was placed in front of the animal’s mouth, and a drop of sweet water (7%–10% sucrose) was given as the unconditioned stimulus (reward). One of two auditory cues (16 kHz or 6 kHz, as cue A or B, 80 dB SPL at the position of the animal’s head) was presented for 2 s as the conditioned stimulus, and after a delay of 1 s, the reward was delivered at a probability corresponding to the auditory cue. The animals were allowed to lick freely at any time, i.e. no punishment was given in response to lick behavior irrelevant to reward delivery. Intertrial intervals were 8 ± 2 s. The reward probability was set at 0.7 (70%) for cue A and 0.3 (30%) for cue B. Each block of 20 trials consisted of: (i) cue A with reward (7 trials); (ii) cue A without reward (3 trials); (iii) cue B with reward (3 trials); (iv) cue B without reward (7 trials). The order of the trials was shuffled for every other block.

During auditory stimulation experiments, the animals were presented with 0.5-s sinusoidal tone stimuli. The experiments proceeded in blocks: each block of stimuli consisted of sequential presentations of 1, 2, 4, 6, 8, 12, 16, and 24 kHz tones (80 dB SPL at the position of animal’s head) at 2 ± 1 s intervals, followed by a delivery of a fixed amount of reward. The order of the sinusoidal tone stimuli within each block was pseudo-randomized. Typically, each imaging session contained 15–20 blocks of stimuli.

The behavioral tasks were controlled and monitored at a 1-kHz sampling rate for training and at 10 kHz for the imaging experiment, by a NIDAQ (USB-6363) and LabVIEW software (National Instruments, Austin, TX). Stimulus sounds (sinusoidal tones) were generated as analog waveforms by the NI-DAQ and a custom-written LabVIEW program. The sinusoidal tone waveforms and the white noise outputs were mixed using an audio mixer, and were delivered from the speaker through a custom-made audio amplifier.

### Wide-field one-photon Ca^2+^ imaging

Wide-field one-photon Ca^2+^ imaging was conducted with a custom-made zoom-variable microscope equipped with an air objective lens (Plan-NEOFLUAR Z 1.0 ×; numerical aperture 0.25; Carl Zeiss, Oberkochen, Germany). The FOV size was set to fit the full extents of the dorsal and temporal glass windows. A set of 470- and 405-nm LEDs (M470L5 and M405L4; Thorlabs, Newton, NJ) and a filter set (MDF-GFP, Thorlabs) were used for the imaging experiments. We used the same set of the dichroic mirror and emission filter for the two excitation lights. An illumination light at 405 nm was used to detect an isosbestic fluorescence point (Allen et al., 2017; Musall et al., 2019) for the correction of hemodynamic contamination. The excitation light intensity under the objective was set to 9.0 mW for the 470-nm light and 5.0 mW for the 405-nm light.

A scientific CMOS (sCMOS) camera (ORCA-Fusion; Hamamatsu Photonics, Shizuoka, Japan) with a resolution of 1024 × 1024 pixels (15.8 μm/pixel) was used as a photodetector. Images were acquired at a frame rate of 40 Hz. The excitation light was switched from frame to frame with a micro-controller (Arduino Mini) and a custom script, resulting in a 20-Hz frame rate for each excitation light. Each LED illumination and frame acquisition timing from the sCMOS camera were input to the DAQ device performing the behavioral task control and analog data acquisition. Note that at the time of imaging, the two glass windows were not positioned in parallel, and there was a difference in their angles (Marmoset 1, ∼20 degrees; Marmoset 2, <5 degrees). We were thus unable to align the optical axis precisely perpendicular to both of the glass windows at the same time. Nevertheless, we believe that this issue had little impact on our results, because we observed sharp boundaries of neocortical vasculature and robust area-specific calcium responses.

### Behavioral tracking

To monitor body movements of the animals, three high-speed CMOS cameras (DMK33UP1300, Imaging Source, Bremen, Germany) and IC Capture acquisition software (Imaging Source) were used. The three cameras recorded the animals’ eye, face, and left-side upper/lower body at sampling rates of 200, 60, and 30 frames/s, respectively. An objective lens (f = 35 mm, MVL35M23 and f = 12 mm, MVL12M23, Thorlabs; f = 3 mm, 89341 Edmund optics, Barrington, NJ) with a focal length adjuster ring (CMV05, Thorlabs) were used to obtain an appropriate FOV from each camera. The custom-written LabView behavioral-task program generated and recorded frame triggers for the three cameras simultaneously. To obtain a uniformly illuminated image from each camera, the array of 56 LEDs (940 nm, OSI5LA5453B; OptoSupply, Hong Kong, China) was mounted at the back of each camera.

In addition to the lick detector, a three-axis accelerometer (KXTC9-2050, Kionix, Ithaca, NY) was affixed to the footrest to monitor hindlimb activity. Signals from these electric sensors were recorded using the Lab-View behavioral-task programs.

### Data acquisition

The signals recorded by the LabView behavioral-task program included: (i) periods and frequencies of sinusoidal tone presentation; (ii) the TTL signals reported by the lick detector circuit; (iii) the signals generated by the accelerometer affixed to the footrest; (iv) the frame acquisition clocks (30, 60, and 200 Hz) for behavioral videography; and (v) the timings of 470-nm and 405-nm LED flashes, corresponding to the timing of frame acquisition for wide-field calcium imaging. For wide-field calcium imaging, HCImage software (Hamamatsu Photonics) was used to grab and store frames. The data were exported as TIFF files for later analysis. IC Imaging Control software (Imaging Source) was used to acquire behavioral videography. The resulting videos were recorded in motion-JPEG format and subsequently exported to MPEG4 format.

### Preprocessing of wide-field imaging data

The original imaging data consisted of frames of emission signals resulting from the alternating 405-nm and 470-nm excitation lights. These two channels were split into separate TIFF files and then processed using a pipeline including motion correction, temporal down-sampling, and baseline estimation. First, motion correction was performed based on the NoRMCorre algorithm (Pnevmatikakis and Giovannucci, 2017) to create pixel-by-pixel correspondence across imaging sessions from the same animal. The frames were then spatially down-sampled four-fold, from 1024 × 1024 to 256 × 256 pixels. The fluorescence baseline was estimated by taking the eighth percentiles of 30-s sliding windows positioned every 15 seconds. The resulting baseline levels were interpolated and smoothed (30-s sliding window averaging) before being used to calculate the Δ*F*/*F* values of the signals.

Hemodynamics correction of the 470 nm-excited signals was based on a previous report (Peters et al., 2021) with slight modifications. The 405 nm- and 470 nm-excited signals from each pixel (in the 256 × 256 pixel frame) were bandpass-filtered using the 4–7 Hz frequency range, which corresponds to the range of the heart rate of awake marmosets (Murphy et al., 2019). Linear regression was performed using the bandpass-filtered signals, to estimate the pixel-by-pixel contribution of the 405 nm-excited (hemodynamic) signals to the 470 nm-excited signals. Finally, hemodynamics-corrected GCaMP signals for each pixel were computed by subtracting the 405 nm-excited signals, multiplied by the regression coefficient computed earlier, from the 470 nm-excited signals.

To alleviate the possible effects of acquisition noise, we applied two-dimensional gaussian filter (1-pixel standard deviation), followed by pixel-by-pixel zero-phase temporal low-pass filter (4.5 Hz cutoff, second-order butterworth filters applied from both directions) to the imaging data, before being used for the later analysis.

### Preprocessing of behavioral data

The output of the lick detector, sampled at 10 kHz, was binarized to ON or OFF at a threshold of 2.2 V. Each ON period was considered to be a raw lick event. Neighboring raw lick events with an interval shorter than 30 ms or duration shorter than 50 ms were considered to be due to “bouncing” and were merged into a single event. The resulting events were removed if they had durations longer than 350 ms, as they were likely to be due to the animal’s hand holding the lick spout. The onsets of the remaining lick events were used as the timings of individual licks.

For the output of the accelerometer on the footrest sampled at 10 kHz, the first derivative of the signal was initially computed to remove any slow offset drift. The root mean-square (RMS) of the signal was then estimated by computing its 100-ms sliding window standard deviation.

DeepLabCut version 2.1 (Mathis et al., 2018; Nath et al., 2019) was used to detect the animals’ behavior from the videography. For each type of key-point estimation, a single model was trained and used for all the behavioral videos of the two animals. In general, estimations with a likelihood lower than 0.9999 were dropped and the corresponding data points were treated as “missing”. The datasets were prepared using the same sampling rate as their source videos (i.e. eye, 200 Hz; hand, 30 Hz; face, 60 Hz) and were stored for later analysis.

For detection of eye motion, a single DeepLabCut model was trained on 161 annotated images from 10 videos. An ellipse was fitted manually to the pupil, and 36 distinct key-points were annotated along its perimeter for use as the key-points to be estimated. The images were subjected to vigorous augmentation procedures: we generated 20 images for each annotated frame using the Python imgaug module. After the DeepLabCut model-based inference step, an ellipse based on the estimated 36 landmarks for each video frame was recovered, by using a least-squares approach. We defined the center of the ellipse as the position of the eye, and the diameter of the pupil was computed as double the length of the ellipse’s semi-major axis. To remove any erroneous estimation (typically due to occlusion of the eye by a finger or blinking activity), each time point was used only when the x-(horizontal-) and y-(vertical-) coordinates of the eye, as well as the pupil diameter, all lay within the 1–99 percentile range. From these data (horizontal/vertical/diameter, in pixels, at 200 Hz), the per-frame motion of the eye (dHorizontal and dVertical, in pixels/frame) and the per-frame net speed of the eye (V, in pixels/frame, non-negative) were also computed.

For detection of hand motion, a single DeepLabCut model was trained on 126 annotated images from seven videos. A bounding rectangle was set around the left hand of the animal, and its geometrical center was annotated as the position of the hand. The images were subjected to vigorous augmentation procedures, and 20 images were generated for each annotated frame using imgaug. After estimation of the x-(horizontal-) and y-(vertical-) coordinates of the hand position at each frame (Horizontal and Vertical, in pixels, at 30 Hz), the per-frame motion of the hand (dHorizontal and dVertical, in pixels/frame) and the net speed (V, in pixels/frame, non-negative) were computed.

To detect face motion, we fine-tuned the “primate-face” model (Clair Witham, Centre for Macaques, MRC Harwell, UK) in DeepLabCut Model-Zoo. The landmarks were adapted for marmoset faces under our experimental rig. It should be noted that our annotations for the ears (key-points #2–7, #20–25 of the original model) and the indents (key-points #11 and #17) did not correspond to the original annotation scheme. In addition, because of limitations with our dataset, the cheek (key-points #14) and the lip (key-points #51–54) were poorly annotated, and the “neck nape” (landmark #55) was not annotated at all. We annotated 36 images extracted from six previously published videos (Correia-Caeiro et al., 2022) and 60 images from six videos of the two animals used in this study. Note that image augmentation was not used for the training of this model. Furthermore, because the area around the animal’s mouth and lower jaw was often masked by the lick spout, only the key-points at or above the level of cheek were used for analysis. For each video (i.e., for each imaging session), two-dimensional deviations from the session-mean position were computed for each key-points at each frame. Face motion components were orthogonalized using principal component analysis (PCA). The scores of the first 18 PCs, which explained 94.17% ± 0.54% (mean ± SD, Marmoset 1, *n* = 10 sessions) and 94.03% ± 0.45% (Marmoset 2, *n* = 9 sessions) of the variance of the total deviations, were stored and used in later analysis.

### Computing class-balanced averages

The trials of our classical conditioning task consisted of 2 × 2 conditions: cue A vs. cue B trials and rewarded vs. non-rewarded trials. These conditions were marginally balanced but conditionally unbalanced; for example, cue A trials consisted of ∼70% rewarded trials and ∼30% non-rewarded trials, whereas cue B trials included ∼30% rewarded and ∼70% non-rewarded trials. Simply taking averages of these unbalanced trials could lead to unwanted overrepresentation of the majority (e.g., rewarded trials in cue A trials) and underrepresentation of the minority (e.g., non-rewarded trials in cue A trials). To alleviate this issue, we performed a balancing procedure, using inversely weighted averaging to compute class-balanced means. For each trial-aligned time point *t*, we have figures of four different conditions: (i) cue A rewarded trials, with a trial number of *N*_A_^*+*^ and mean value of *M*_A_^*+*^(*t*); (ii) cue A non-rewarded trials, *N*_A_^*−*^ and *M*_A_^*−*^(*t*); (iii) cue B rewarded trials, *N*_B_^*+*^ and *M*_B_^*+*^(*t*); and (iv) cue B non-rewarded trials, *N*_B_^*−*^ and *M*_B_^*-*^(*t*). From these values, the balanced averages for cue A and cue B trials, *M*_A_(*t*) and *M*_B_(*t*), were computed as:

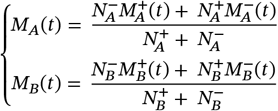

### Encoding models

We used linear encoding models with ridge regression (Karabatsos, 2018; Musall et al., 2019) with slight modifications. Instead of using trial-aligned peri-event time histograms (PETHs), we used one whole imaging session over its full duration (i.e., including both trial time and inter-trial intervals) to estimate and validate a single set of encoding models.

The task-related columns of the design matrix included cue-related and reward-related variables. Cue A trials and cue B trials were encoded separately as cue-related variables. Encoding of trials included time-to-time bias terms; individual timepoints in the trial (i.e. frames 1, 2, …, 160, starting from the tone onset and up to +8 s after the tone onset) were represented as separate columns for each of the tone types in the design matrix. Reward was represented (only in the timepoints corresponding to rewarded trials) as a set of time-to-time bias terms from +3 s to +8 s after the tone onset. Together, the diagonal representations of cue A trials, cue B trials, and rewards comprised the task-related columns of the design matrix.

The motion-related columns of the design matrix included: Lick (lick counts), Hand/Foot (hand Horizontal, hand Vertical, hand dHorizontal, hand dVertical, hand V, foot sensor RMS), Eye/Pupil (eye Horizontal, eye Vertical, eye dHorizontal, eye dVertical, eye V, pupil diameter), and Face (face deviation PC scores). These motion-related variables were resampled to the 20 Hz imaging frame rate. Because the face PC scores presumably contained a non-negligible amount of information overlapping with the other behavior variables, they were decorrelated from the other behavior variables using QR decomposition before being added to the design matrix. To account for the preparatory and feedback activity corresponding to the behavioral output at each time point, time-shifted versions (from –20 frames to +20 frames, 41 columns in total per variable) of each behavior-related variable were also included.

Pixel calcium activity was normalized by computing the Z-scores. To speed up calculation of the encoding models, singular value decomposition (SVD) was applied, and the first 8500 components from the largest, corresponding to >99% of the total variance, were used for the analysis. On the side of the design matrix, all columns were Z-scored prior to the analysis. A linear model was fitted by ridge regression, to predict each singular-value component (i.e. each row vector of the product of the right singular matrix and the singular value matrix) based on the design matrix of the session. The pixel-by-pixel prediction in Δ*F*/*F* values was subsequently recovered by multiplying with the spatial components (i.e. the left singular matrix), and by referring to the mean and standard-deviation values of each pixel.

Ten-fold cross validation was used: 90% of the timepoints were used to estimate a linear model including its intercept term, and the fitting performance of each model was validated by computing the coefficient of determination for the remaining 10% of the timepoints (cross-validation R^2^, or cvR^2^). Throughout this study, out of the 10% of timepoints, only those timepoints corresponding to trials (i.e., where the tone-related variables had non-zero values) were used to compute cvR^2^ values. To compute the unique contribution of a specific set of variables, the columns corresponding to the variables of interest were first temporally shuffled, then the reduction in cvR^2^ values in the resulting linear model (ΔcvR^2^) was calculated. Each ΔcvR^2^ value was rectified to ensure that it had a non-negative value. To alleviate sampling bias, five different splits of ten-fold cross validation were performed, and their mean value was reported.

When analyzing the responses of the auditory stimulation experiments, the unique contributions of cue A-, cue B-, and non-cue tone-related variables were computed for the 3-s time window after stimulus onset. When computing the cvR^2^ values, imbalances in the number of trials were taken into account by using only those trials and timepoints where the tones of concern were presented; given *N*_A_, *N*_B_, and *N*_N_ as the number of cue A, cue B, and non-cue-tone trials, respectively, the cvR^2^ values were computed separately for cue A-, cue B-, and non-cue tone-related variables by using the corresponding *N*_A_ × 3 s, *N*_B_ × 3 s, or *N*_N_ × 3 s after the stimulus onset of the corresponding trials.

### Hierarchical clustering

For the clustering of calcium activity, trial-aligned PETHs were generated for the 12 ROIs. The 1-s window before tone onset was considered as the baseline, and the mean activity during this period was subtracted for each trial. The signals that lay within the 6-s window from the tone onset (i.e., until 3 s after the reward timing) were used as the within-trial signals, whereas those within the –4 to –1 s and +6 to +9 s periods were considered to be pre-trial and post-trial signals, respectively. For clustering of movement-related unique contributions, all the weights assigned to all the movement-related variables were used for each ROI. For each of the three categories of signals (i.e., within-trial activity, out-of-trial activity, and movement-related unique contributions), the values pooled across all the trials from all the imaging sessions were concatenated, and the Pearson’s correlation coefficient, *R*, was computed for each pair of ROIs. Agglomerative (bottom-up) hierarchical clustering was performed using the value (1 – *R*) as the distance metric. Weighted averaging of correlation coefficients was used as the linkage method. The arbitrary threshold *R* = 0.4 was used to define clusters.

### Statistical testing: neural and behavioral activity PETHs

To calculate the PETHs of the calcium activity, the mean of the 1-s pre-cue activity of each trial was considered as the baseline and was subtracted from the subsequent trial-aligned trace. No baseline subtraction was performed for PETHs of behavior-related parameters.

For the results of the classical conditioning task, class-balanced within-session average traces were computed (Marmoset 1, *n* = 10 sessions; Marmoset 2, *n* = 9 sessions). The Wilcoxon signed-rank test was performed for each time point along the trial to compare the two trial conditions (cue A vs. cue B, or rewarded vs non-rewarded). The significance level threshold was set to α = 0.05.

### Statistical testing: ROI-by-ROI comparison

A resampling-based testing (Russo et al., 2020; Talluri et al., 2023) was used to test the statistical significance of ROI-based values. For each session, we prepared *N* pairs of values, with *N* being the number of sessions (data vs. chance for single-ROI comparison, data 1 vs. data 2 for between-ROI comparison). To test whether the mean difference between the two sets of values was statistically significant, the chance distribution of the difference was estimated by sampling from the values with replacement, without distinction between the two classes under consideration, randomly giving them the class annotations and computing the mean difference between the two classes. This resampling procedure was performed for 30,000 iterations, and we examined whether the value derived from the original data lay outside of the 95-percentile range of the resampled distribution.

To compute chance-level ΔcvR^2^ values for each session, we used the full-model cvR^2^ values based on the five different iterations (i.e. cross-validation splits). By selecting one of the five values as the reference, we computed the ΔcvR^2^ values of the other four full models, and defined their mean as the chance-level ΔcvR^2^.

To compare values between ROIs, we performed multiple paired resampling tests as described above, with Holm-Bonferroni correction. The significance level threshold was set to α = 0.05.

