## Supplementary Figures 1-7 for "Movement-independent representation of reward-predicting cues in the medial part of the primate premotor cortex"

bioRχiv

### Supplementary Figures 1–7

### Movement-independent representation of reward-predicting cues in the medial part of the primate premotor cortex

Keisuke Sehara, Masashi Kondo, Yuka Hirayama, Teppei Ebina, Masafumi Takaji, Akiya Watakabe, Ken-ichi Inoue, Masahiko Takada, Tetsuo Yamamori, and Masanori Matsuzaki

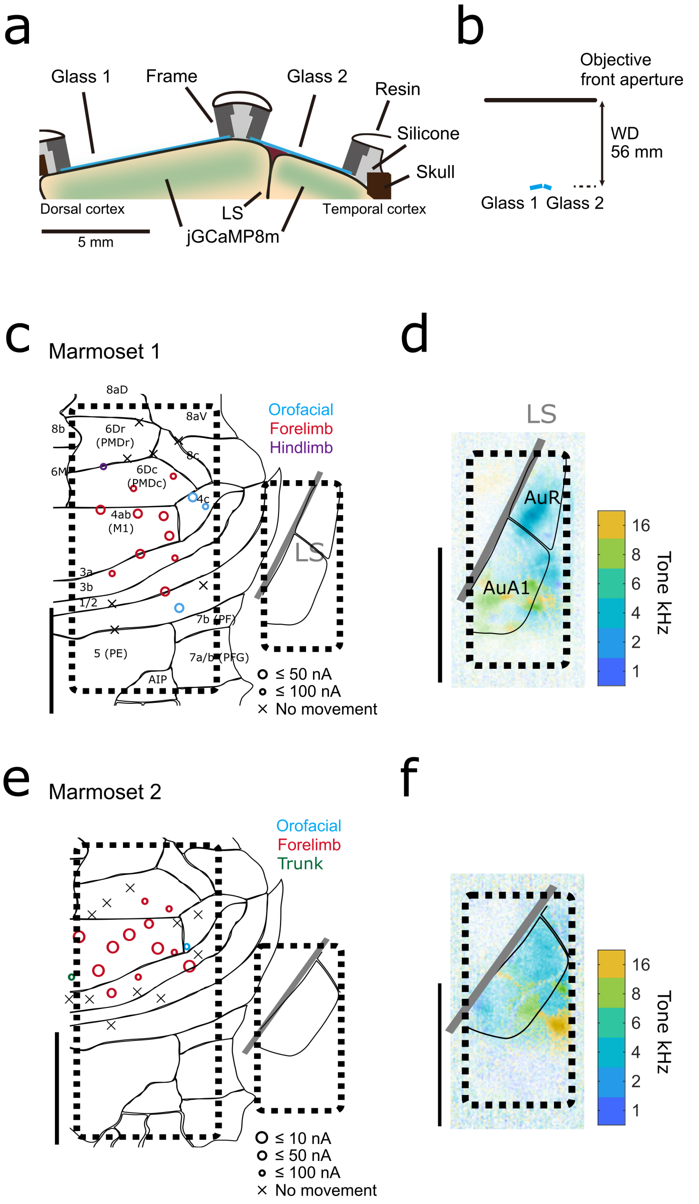


**Supplementary Figure 1: Annotation of cortical areas.**

**a** Schematic illustration of chronic window implants on the cortical surface. One glass coverslip (Glass 1) was placed over the dorsal cortex, and the other one (Glass 2) was over the lateral side of the somatosensory cortex and the superior temporal cortex. These glass windows were secured by frames. **b** A schematic illustrating the arrangement of the imaging windows and the objective lens exit. WD, working distance. **c,e** The results of the ICMS experiment on Marmoset 1 (**c**) and Marmoset 2 (**e**). The type and the size of the marker indicates the response threshold, and the color represents the type of the evoked movement. The standard brain atlas (Paxinos et al., 2012) is overlaid in accordance with the determined PMdc-M1 border (for the main window). Scale bar, 5 mm. **d,f** Tonotopy mapping of the second imaging window in Marmoset 1 (**d**) and Marmoset 2 (**f**). The border between areas rA1 and cA1 was determined according to the area responding to low-frequency tones. Scale bar, 5 mm.

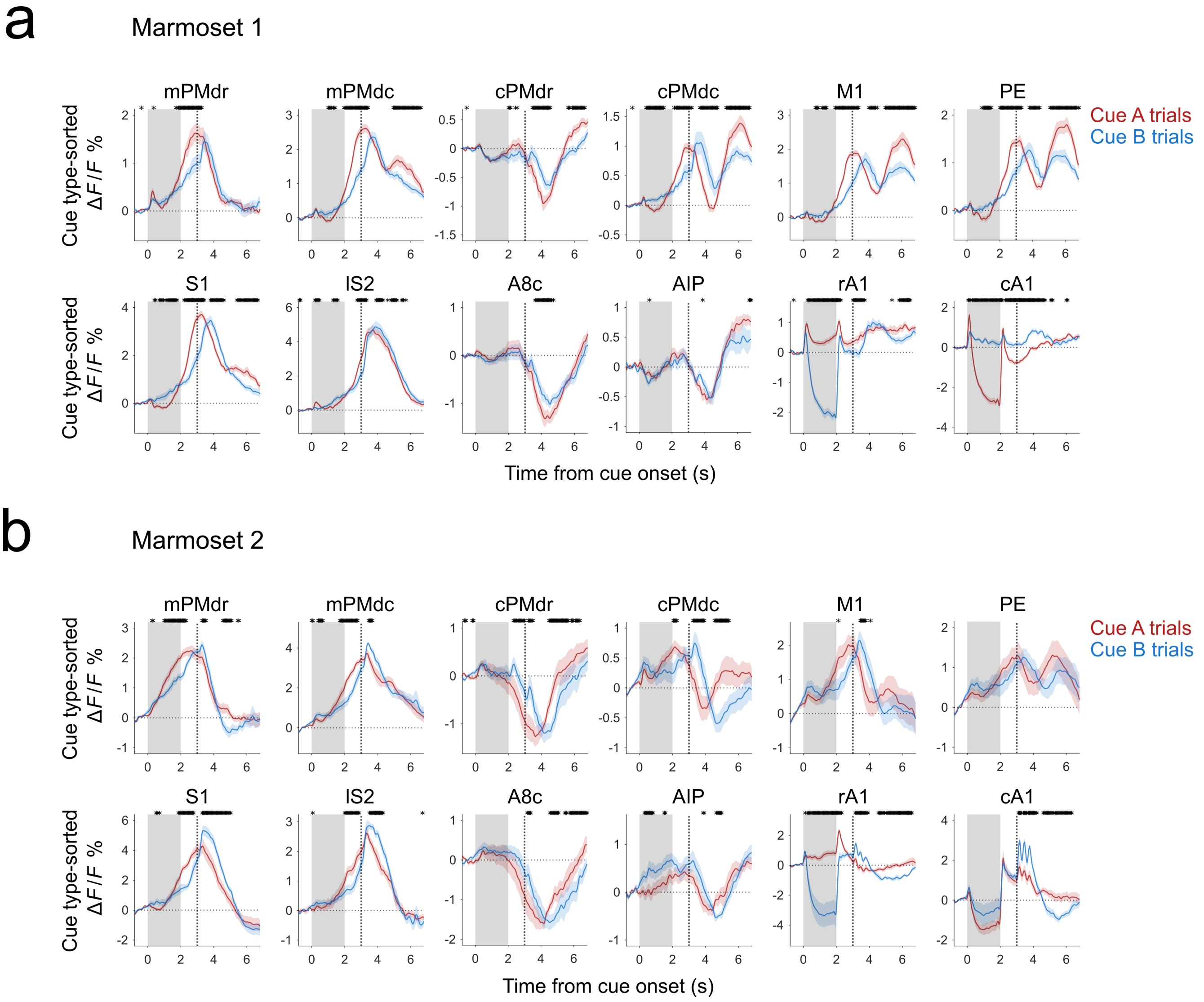


**Supplementary Figure 2: ROI-based analysis of cue-aligned calcium responses.**

**a,b** Responses of the 12 ROIs to conditioned stimuli, cue A (red) and cue B (blue), in Marmoset 1 (**a**) and Marmoset 2 (**b**). Session-averaged baseline-subtracted traces were computed as weighted averages to account for the imbalance in numbers between rewarded and non-rewarded trials. The mean and SEM of the session-mean traces are shown. Gray shaded areas represent the duration of cue presentation, and black dotted lines indicate the reward timing. $^{*}P < 0.05$, Wilcoxon signed-rank test between cue A and cue B trials. Marmoset 1, $n = 10$ sessions; Marmoset 2, $n = 9$ sessions.

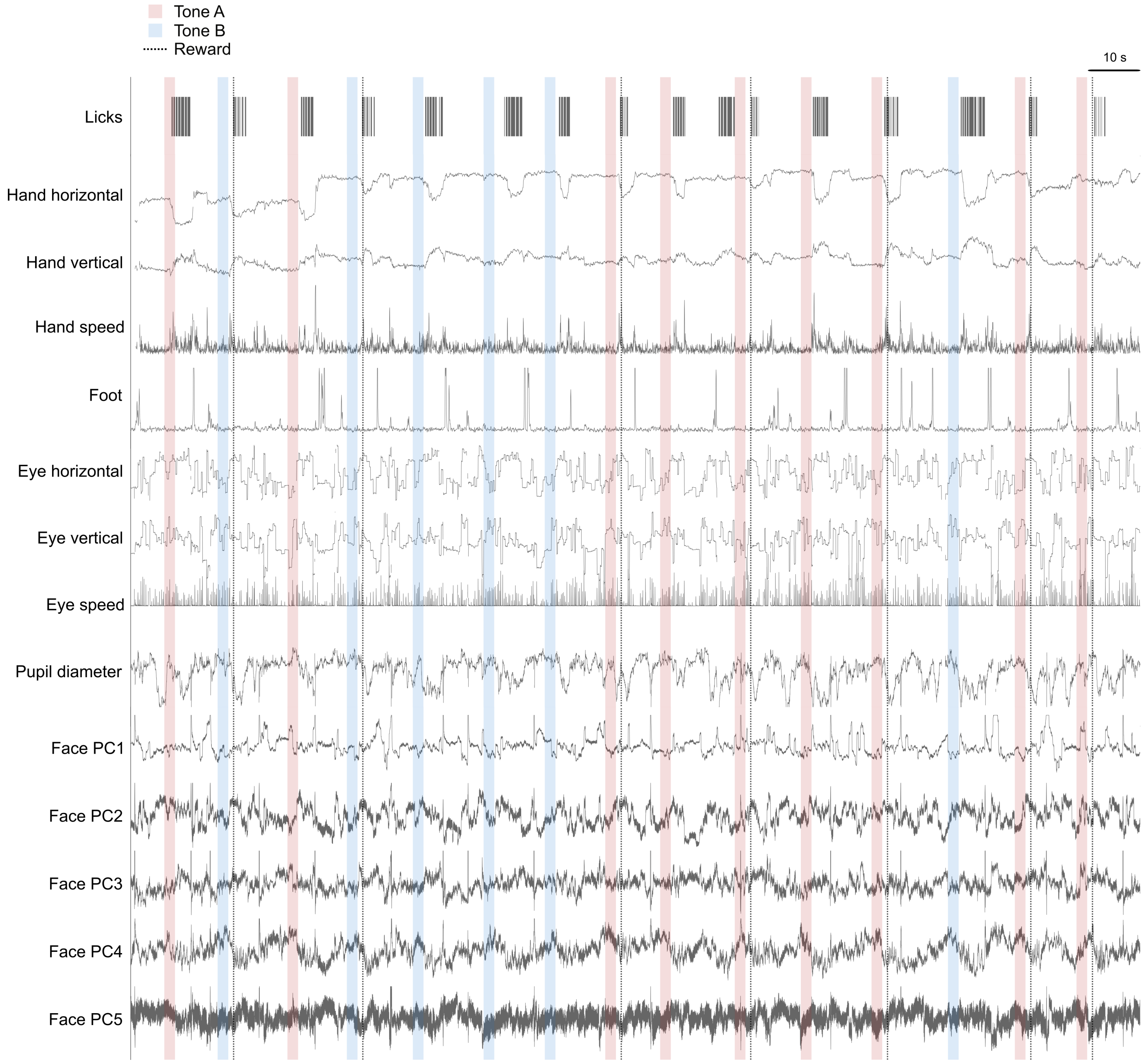


**Supplementary Figure 3: Single-trial dynamics of behavioral parameters.**
Representative traces of behavioral variables in Marmoset 1. Shaded areas and dotted lines represent cue presentations and reward delivery timings. Some traces were clipped above arbitrary thresholds.

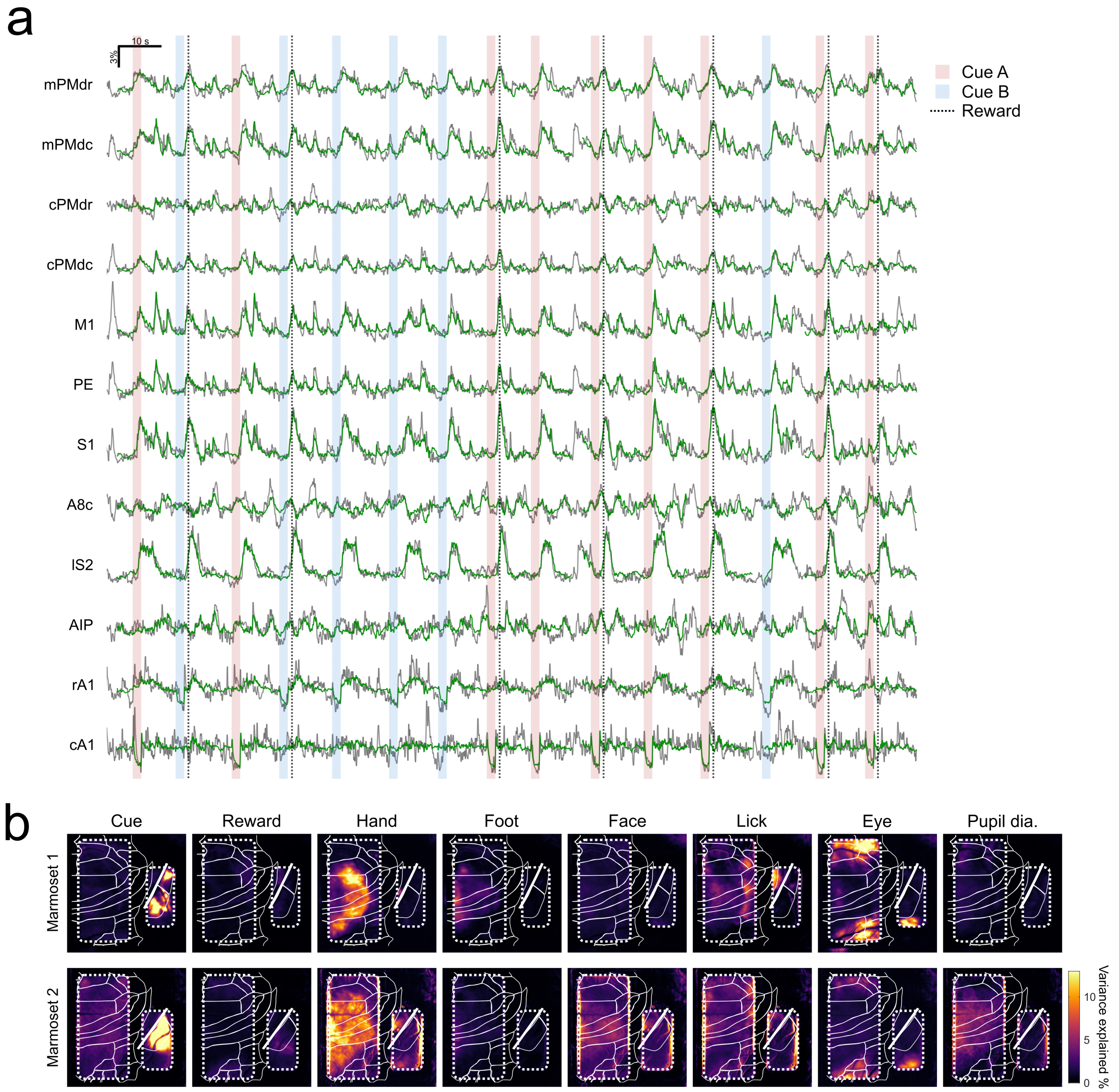


**Supplementary Figure 4: Performance of the ridge linear regression models.**

**a** Representative traces of recorded (gray) and model-predicted (green) calcium responses of 12 ROIs during the session shown in Supplementary Fig. 3. Shaded areas represent cue presentation (cue A, red; cue B, blue), and dotted lines represent reward delivery. Predicted calcium responses are missing in some timepoints due to failure in the prediction of movement-related variables. Note that there were four 2-s time windows without predicted responses. This was because a single missing movement-related value for each window resulted in a prediction failure for two seconds because of the convolution procedure, i.e., linear summation of time-shifted series of values, of the linear encoding models. **b** Unique contributions of distinct sets of variables to the performance of the encoding models. The average pixel-by-pixel explained variances ($\Delta cvR^2$) across sessions are color-coded for each animal. Marmoset 1, $n = 10$ sessions; Marmoset 2, $n = 9$ sessions.

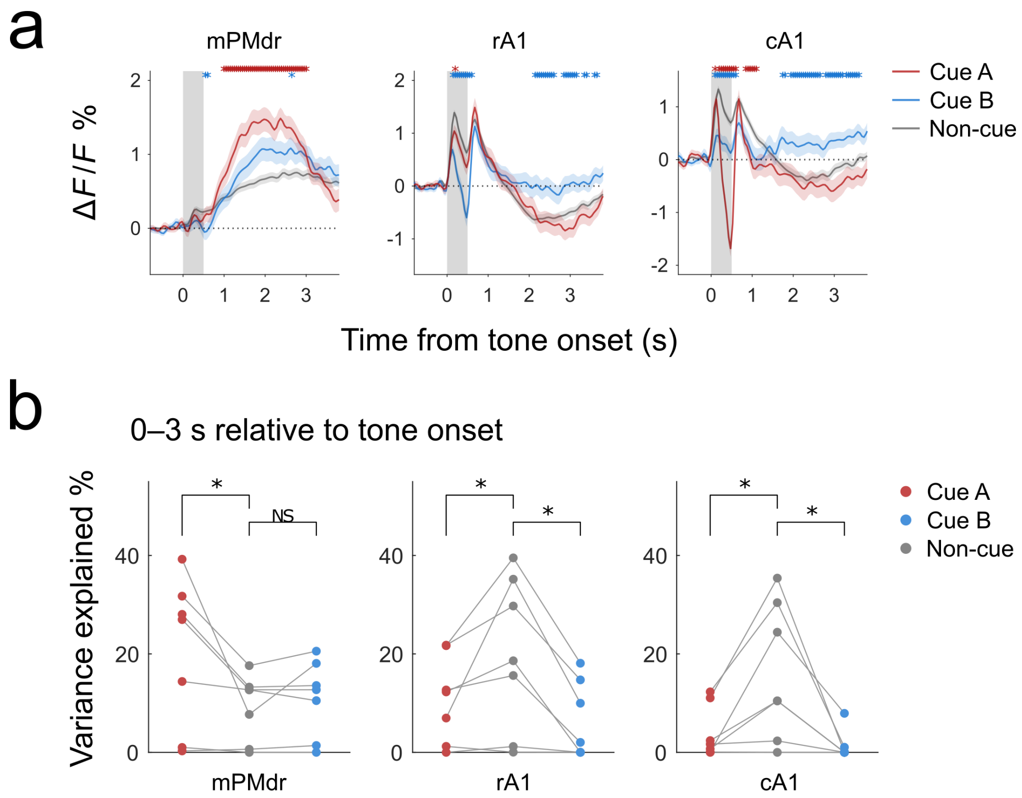


**Supplementary Figure 5: Responses to cue and non-cue tones during auditory stimulation experiments.**
**a** Baseline-corrected calcium responses of mPMdr, rA1, and cA1 to 500-ms stimuli of different tone frequencies (1, 2, 4, 6, 8, 16, and 24 kHz). Mean and SEM of the responses in the pooled 880 trials from two animals are shown. Cue A tone, $n = 46$ trials from 3 sessions (Marmoset 1) and 61 trials from 4 sessions (Marmoset 2); Cue B tone, $n = 47$ (Marmoset 1) and 62 (Marmoset 2) trials; Non-cue tones, $n = 233$ (Marmoset 1) and 313 (Marmoset 2) trials. $^*P < 0.05$, Wilcoxon signed-rank test with Holm–Bonferroni correction, Cue A tone vs. non-cue tones (red), and Cue B tone vs. non-cue tone (blue). **b** Linear encoding models were constructed in a similar manner as in the classical conditioning task. Unique contributions ($\Delta cvR^2$ values) in mPMdr, rA1, and cA1 were computed during the 3-s period after the stimulus onset. The $\Delta cvR^2$ values were compared by testing the differences between either cue A and non-cue tones, or cue B and non-cue tones. $^*P < 0.05$, pairwise resample tests with Holm-Bonferroni correction. $n = 7$ sessions ($n = 3$ from Marmoset 1, and $n = 4$ from Marmoset 2).

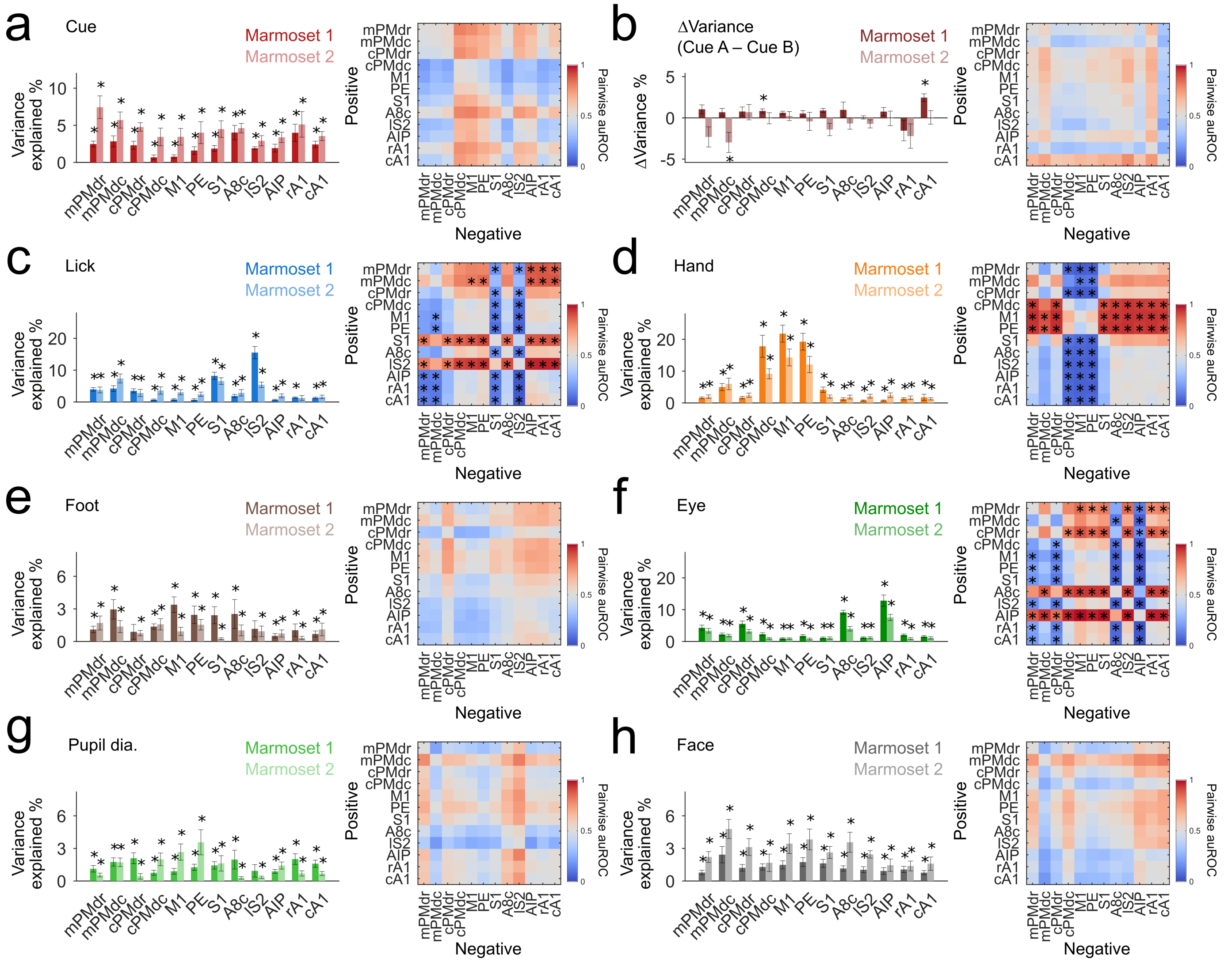


**Supplementary Figure 6: Unique contributions of distinct sets of variables during the response period.**
**a** Left, unique contributions of the cue variables during the response period for each ROI, based on the values of $n = 10$ (Marmoset 1) and $n = 9$ (Marmoset 2) sessions. Right, matrix of pairwise auROC values, based on the values of $n = 19$ pooled sessions from two animals. $^*P < 0.05$, resample test (left; comparison with the chance levels), and resample test with Holm-Bonferroni correction (right). **b** Similar to **a** but for the differences between cue A-related and cue B-related unique contributions, ΔVariance(Cue A – Cue B), along the procession of the trial. **c–h** Left, unique contributions of the six movement-related variables: lick (**c**), hand (**d**), foot (**e**), eye (**f**), pupil diameter (**g**), and face (**h**) during the response period for each ROI. Right, matrix of pairwise auROC values. The values are color-coded. $^*P < 0.05$, resample test (left), and resample test with Holm-Bonferroni correction (right).

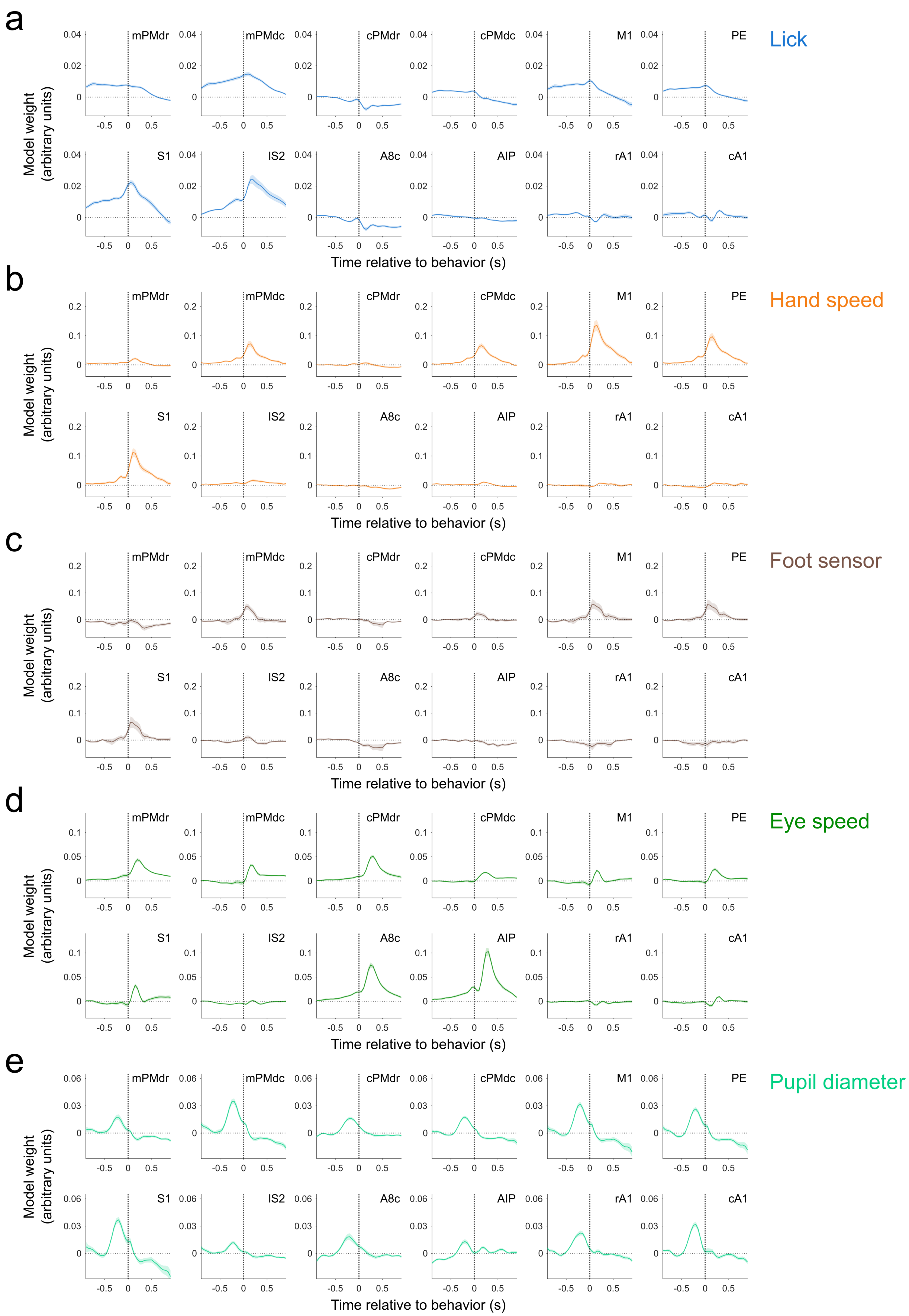


**Supplementary Figure 7: Full-model weights attributed to representative movement-related variables.**
**a-e** Model weights of lick (**a**), hand speed (**b**), foot (**c**), eye speed (**d**), and pupil diameter (**e**) for the ±1-s time window. The mean and the SEM values of all the pooled sessions from the two animals are plotted against the time relative to the timing of each behavior under consideration. $n = 19$ sessions from the two animals ($n = 10$ from Marmoset 1, and $n = 9$ from Marmoset 2).
